# MetaReact: A Reaction-Aware Transformer for End-to-End Prediction of Drug Metabolism

**DOI:** 10.64898/2026.03.14.711529

**Authors:** Yitian Wang, Jingxin Rao, Wei Zhang, Yuqi Shi, Chuanlong Zeng, Rongrong Cui, Yinquan Wang, Jiacheng Xiong, Xutong Li, Mingyue Zheng

## Abstract

Accurate prediction of drug metabolites and enzyme selectivity is essential for rational drug design and safety assessment. However, existing computational approaches are often limited to specific enzyme families or reaction types, lacking the capacity to model enzyme-subtype specificity and prioritize major metabolites. Here, we present MetaReact, an end-to-end generalizable Transformer-based model that unifies the prediction of metabolic enzymes, metabolites, and sites of metabolism (SOM). By integrating structure-aware encoding ReactSeq, a chemistry reaction-based pretraining, MetaReact consistently outperforms state-of-the-art methods across multiple benchmarks under three settings: enzyme-agnostic, enzyme-completion, enzyme-conditioned. Notably, it achieves 60% Top-3 accuracy in identifying major metabolites and superior CYP450 enzyme-subtype prediction/SOM recognition. Case studies validate its applicability to complex natural products, synthetic cannabinoids, and clinical candidates, facilitating toxicity assessment and molecular optimization. This scalable, rule-free solution advances human metabolism modeling, with potential for computational pharmacokinetics and early drug discovery.

## 1 Introduction

Drug metabolism is a pivotal determinant of pharmaceutical efficacy and safety, as enzymatic conversion of drug molecules into metabolites often alters their physicochemical characteristics, pharmacological activity, or potential toxicity—key factors linked to late-stage drug development failures.^1–3^ Typically proceeding via Phase I functionalization (e.g., oxidation, reduction) followed by Phase II conjugation (with endogenous groups to enhance solubility and excretion efficiency)^4^, such metabolic processes can introduce unforeseen safety risks even for structurally related metabolites^5^. Consequently, early and accurate prediction of drug metabolites is therefore critical for informed decision-making in pharmaceutical development.^6,7^ Regulatory guidelines (e.g., U.S. Food and Drug Administration [FDA] and International Council for Harmonisation of Technical Requirements for Pharmaceuticals for Human Use [ICH] M3 (R2)) further emphasize identifying and characterizing metabolites accounting for >10% of total drug-related exposure before Phase III clinical trials.^8–10^

Common preclinical approaches for metabolic profiling include recombinant enzyme systems, human liver microsomes (HLMs), primary human hepatocytes, and animal models.^7,11^ While *in vitro* systems are cost-effective, they often fail to recapitulate the full complexity of metabolism in living organisms.^12–14^ For example, while recombinant enzyme incubations can confirm the involvement of specific enzymes, they lack physiological context. HLMs, while widely used, primarily comprise cytochrome P450 enzymes (CYPs), uridine 5’-diphospho-glucuronosyltransferases (UGTs), and esterases, and thus fail to capture the activity of other metabolic enzymes.^15,16^ Animal models offer closer physiological relevance but are limited by interspecies differences.^17–19^ Humanized chimeric mice with livers repopulated by >80% human hepatocytes have recently shown promise in better recapitulating human metabolic pathways.^20–22^ However, their widespread adoption in routine screening remains limited by high costs and restricted availability.

Computational approaches enable rapid, large-scale screening of candidate compounds during early-stage drug development. Current computational strategies for predicting drug metabolism generally fall into two categories. The first focuses on identifying sites of metabolism (SOMs)—specific atoms or bonds within a molecule that are vulnerable to enzymatic modification. Representative models include RS-Predictor^23^, SOMP^24^, SMARTCyp3.0^25^, FAME3^26^, Rainbow XenoSite^27^, GNN-SOM^28^ and GLMCyp^29^. Many rely on precomputed features such as density functional theory (DFT) activation energies, topological descriptors, and steric accessibility, and are largely limited to CYP450-mediated reactions. GNN-SOM overcomes this limitation by leveraging an end-to-end graph neural network (GNN) to learn atomic representations from molecular graphs and predict the likelihood of metabolism at specific atoms or bonds, conditioned on two-digit EC number.^28^ GLMCyp is a deep learning model for predicting CYP450 reaction sites on small molecules. By combining 2D molecular graph features, 3D structural representations from Uni-Mol^30^, and CYP450 protein features from ESM-2^31^, it accurately identifies SOMs for nine human CYP450 isoforms.^29^ While these models outperforms conventional machine learning models, it remains restricted to resolving enzyme subtype specificity, particularly within closely related CYP450 isoforms.

The second category focuses on directly predicting the chemical structures of metabolites generated from the parent compound. Most current tools in this category are rule-based, such as SyGMa^32^, RD-Metabolizer^33^, GLORYx^34^, Metabolic Forest^35^, CyProduct^36^, and BioTransformer3.0^37^. These approaches rely on expert-defined transformation rules that are only activated when the substrate precisely matches a predefined pattern. As a result, the utility of such approaches is constrained by insufficient rule coverage and specificity; moreover, greater rule complexity may further increase the likelihood of false positives. To overcome the shortcomings of rule-based methods, recent advances have introduced template-free deep learning models that learn transformation patterns directly from data. MetaTrans formulates metabolite prediction as a SMILES-to-SMILES translation task, utilizing a Transformer architecture pretrained on general chemical reaction datasets and subsequently fine-tuned on metabolic reaction data.^38^ MetaPredictor integrates SOM tagging with a prompt-based strategy to guide metabolite generation in an end-to-end manner.^39^ While these deep learning models demonstrate improved generalization over rule-based systems, they still lack enzyme-specific guidance and fail to capture the selectivity among enzyme families or subtypes, limiting their adaptability to different needs in early discovery, mechanistic metabolism studies, and medicinal chemistry optimization.

Recent advances like DeepMetab have unified CYP450-mediated metabolism prediction (substrate identification, SOM localization, metabolite generation) via a GNN-based multi-task framework, using rule-based metabolite generation post-SOM prediction to ensure mechanistic consistency.^40^ Yet even these models suffer from limited generalizability, poor enzyme specificity, and inadequate coverage of non-CYP enzymes, creating a critical gap for a unified, flexible modeling framework. Notably, no existing model simultaneously addresses the dual needs of medicinal chemists and pharmacokineticists: predicting enzyme-substrate selectivity across diverse enzyme families, and inferring SOMs and metabolite structures under both known and unknown enzymatic contexts.

To fill this gap, we introduce MetaReact, a deep learning framework designed for comprehensive drug metabolism prediction across diverse enzymatic contexts. By leveraging ReactSeq^41^, a reaction-aware molecular representation that encodes atomic and bond-level changes between reactants and products, MetaReact inherently captures reaction-center information to support SOM identification. Pretrained on a broad collection of general organic reactions and fine-tuned on a large curated dataset of metabolic reactions, MetaReact supports three task settings (*enzyme-agnostic*, *enzyme-completion* and *enzyme-conditioned*) for flexible adaptation to diverse drug discovery use cases. These capabilities enable direct inference of metabolic enzymes, SOMs, and metabolite structures from a given substrate, offering a unified, generalizable tool to advance both medicinal chemistry and pharmacokinetic research.

## 2 Results

### 3.1 Overview of MetaReact

MetaReact comprises a Transformer encoder-decoder architecture (Figure 1A) and employs a reaction-aware molecular representation (ReactSeq) (Figure 1B), forming a unified sequence-to-sequence framework that overcomes the fragmentation and task-specific rigidity of existing metabolism prediction tools. This framework supports three complementary metabolic prediction modes that reflect distinct real-world metabolic analysis scenarios (Figure 1C): **(1) *Enzyme-agnostic* prediction**, which predicts metabolites from substrates alone, enabling early-stage screening where catalyzing enzymes are unknown; **(2) *Enzyme-completion* prediction**, which jointly infers both the metabolizing enzyme (at family and subtype levels) and the corresponding metabolite by appending a placeholder token “<unk>” to the substrate input, enabling the elucidation of unknown enzymatic pathways relevant to drug clearance and toxicity. (**3) *Enzyme-conditioned* prediction**, which predicts metabolites when the catalyzing enzyme is known (e.g., identified through recombinant enzyme assays), uses the concatenated substrate-enzyme pair as input to facilitate targeted molecular optimization aimed at mitigating enzyme-specific metabolic liabilities. Together, they encompass the operational landscape of metabolic prediction, enabling systematic evaluation of model robustness across diverse application contexts and degrees of enzymatic prior information. By accepting different combinations of inputs, such as substrates alone, substrates with unknown enzymes, or substrates with known enzymes, MetaReact can flexibly execute all three prediction modes within the same unified framework.

**Figure 1.**
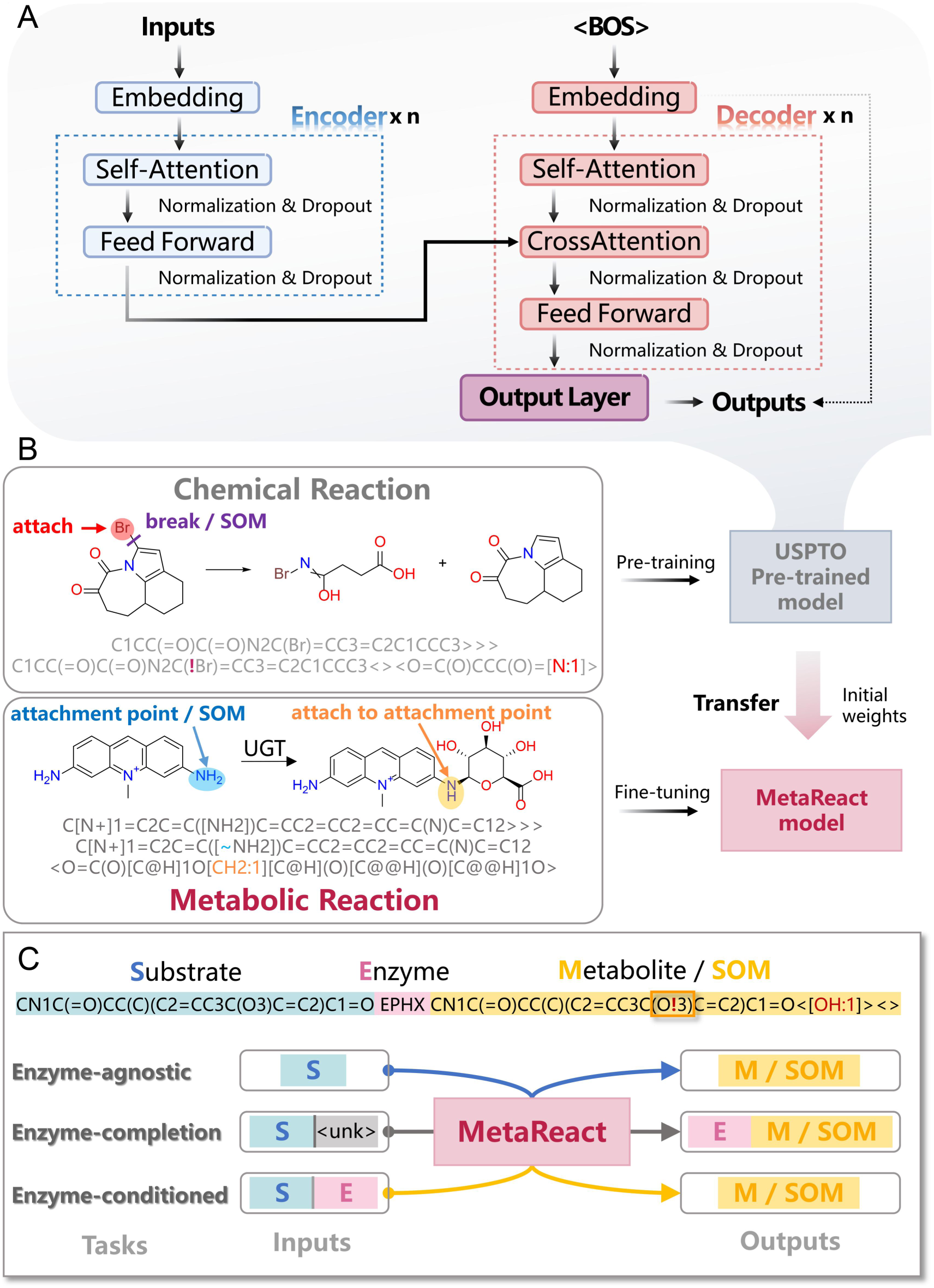
Overview of the MetaReact model. **(A)** MetaReact model architecture. The encoder uses self-attention to generate context-aware representations of the input sequence. The decoder generates output tokens autoregressively, using both self-attention and cross-attention to integrateinformation from the encoder. **(B)** Pretraining on general chemical reactions and fine-tuning on metabolic reactions, both represented using the reaction-aware ReactSeq format that encodes atomic and bond transformations. **(C)** Three task-specific input formats guide the model to predict metabolites from substrates (*enzyme-agnostic*), infer both enzyme and metabolite (*enzyme-completion*), or predict metabolites given an enzyme (*enzyme-conditioned*).

To enhance its performance on these tasks, MetaReact integrates two key components: a reaction-aware molecular representation (ReactSeq) and a two-stage transfer learning pipeline (Figure 1B). ReactSeq explicitly encodes atom- and bond-level transformations in both substrates and products, enabling intrinsic identification of SOMs by highlighting reaction centers. Building on this representation, the two-stage transfer learning pipeline allows MetaReact to acquire both general chemical reactivity knowledge and metabolism-specific insight. The model is first pretrained on 479,035 reactions from the USPTO-MIT database^42^ to learn broadly generalizable reaction patterns, and then fine-tuned on a curated set of 62,695 single-step enzymatic drug metabolism reactions to capture transformations specific to metabolic biotransformations.

### 3.2 E*nzyme-agnostic*: metabolite prediction in the absence of enzyme information

In the *enzyme-agnostic* setting, the model predicts metabolites directly from the substrate without enzyme information. This scenario highly relevant to early drug discovery where catalyzing enzymes are often unknown. We first evaluated MetaReact on the internal test set comprising 6,271 metabolic reactions, which was partitioned according to metabolic pathways to effectively prevent data leakage (for details, see the Methodology section). The model achieved Top-5, Top-10, and Top-20 recall of 60.36%, 70.59%, and 77.89%, respectively (Table 1). In comparison, models using SMILES representations instead of ReactSeq, or models without chemistry reaction-based pretraining, exhibited substantially lower performance, highlighting the contribution of ReactSeq and reaction-aware pretraining to improved metabolic prediction.

**Table 1.** Ablation studies of MetaReact on the internal test set.

| Strategies |  | Metrics |  |  |
| --- | --- | --- | --- | --- |
| ReactSeq | Pretraining | Top-5 Recall | Top-10 Recall | Top-20 Recall |
| ✓ | ✓ | **60.36%** | **70.59%** | **77.89%** |
| ✓ | - | 56.60% | 68.08% | 75.17% |
| - | ✓ | 56.13% | 61.73% | 63.01% |
| - | - | 54.16% | 59.50% | 60.68% |

**Note:** The best results are shown in bold. *ReactSeq* refers to using the ReactSeq representation instead of standard SMILES. Pretraining indicates the use of USPTO-based pretraining. Top-k recall denotes the recall of the top-k predictions made by MetaReact (k = 5, 10, 20). The highest scores are indicated in bold.

We then evaluate the model on the MetaTrans dataset, a benchmark introduced by Litsa et al.^38^, which includes 65 drugs and 179 experimentally validated metabolites. MetaReact consistently outperformed both rule-based methods (SyGMa, GLORYx, and BioTransformer) and Transformer-based baselines (MetaTrans, MetaPredictor). As summarized in Figure 2A-C and Table S1-2, it achieved the highest accuracy across enzyme classes: for oxidase-catalyzed reactions, 90 correct predictions out of 100 at the Top-13 cutoff (vs. 80 for MetaPredictor, 70 for MetaTrans/GLORYx, 68 for SyGMa, and 81 for BioTransformer). In Phase II metabolism (UGTs), it also ranked first (8/17 correct) and maintained strong performance for sulfotransferases (SULTs) and other transferases where rule-based tools performed poorly. For hydrolases, MetaReact, SyGMa, and GLORYx performed comparably, while for reactions with unspecified enzymes, MetaReact retained a clear advantage (Figure 2A, and Table S1). Across coverage metrics (Figure 2B, and Table S2), it recovered at least one correct metabolite in 96.9% of cases, half in 90.8%, and all in 60.0%. Precision-recall analysis (Figure 2C) further confirmed superior performance at multiple cutoffs.

**Figure 2.**
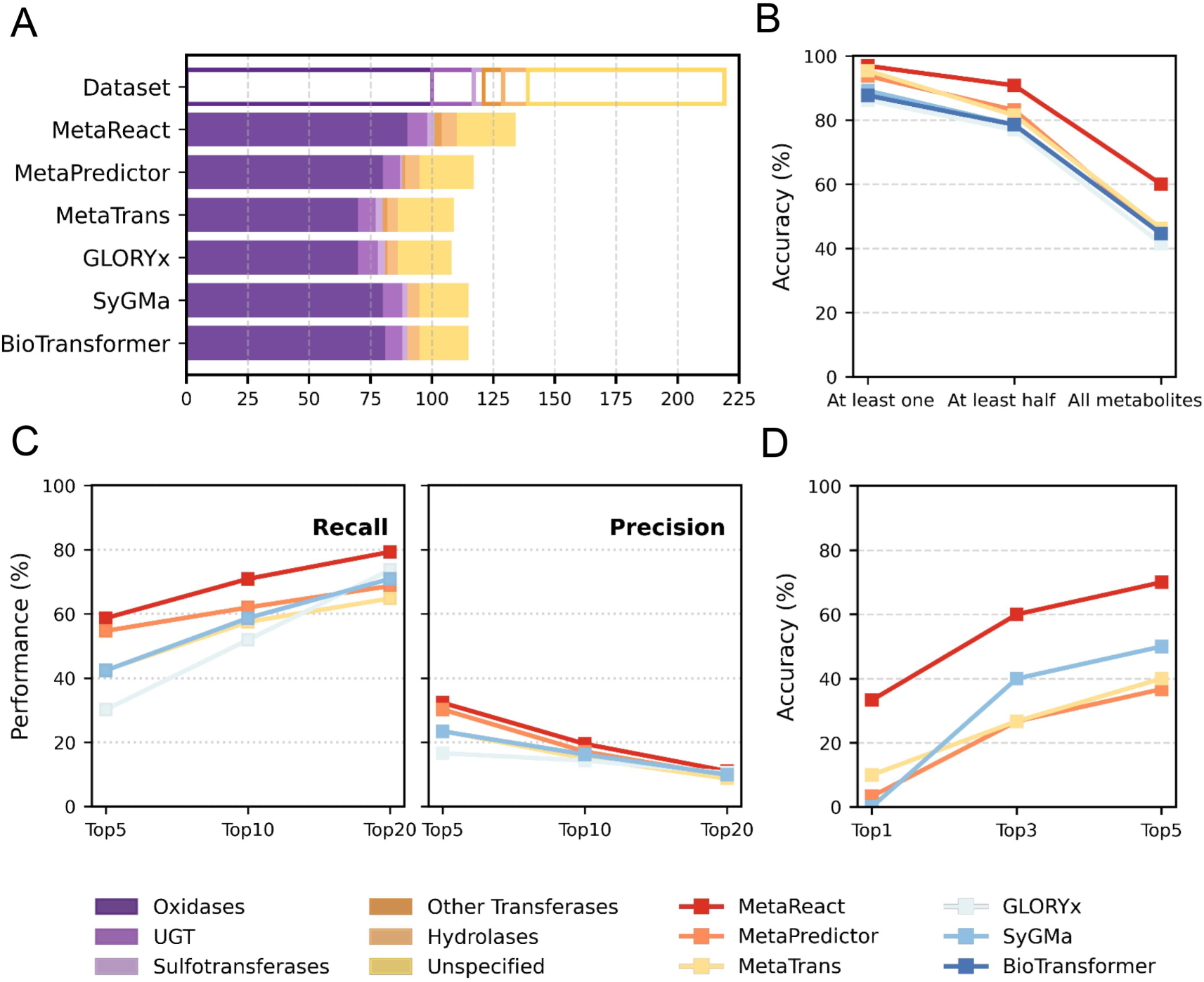
Comparative performance of different metabolite prediction methods in the *enzyme-agnostic* setting. **(A)** Number of correctly identified metabolites for each enzyme family in the MetaTrans dataset, based on the Top-13 predictions per reaction. Bar segments represent different enzyme families. *Dataset* denotes the total number of metabolites for each enzyme family in the dataset. **(B)** Accuracy comparison using three reaction-coverage metrics within the Top-13 predictions. **(C)** Comparison of precision and recall for metabolite identification at Top-5, Top-10, and Top-20 predictions. BioTransformer is excluded here because its output size is fixed at 13 candidates. **(D)** Accuracy for predicting major metabolites on the L-data dataset at Top-1, Top-3, and Top-5 cutoffs.

To specifically assess the model’s ability to predict major metabolites, which are of greater clinical and regulatory significance than general metabolite profiling, we introduced an additional external tests data containing manually curated major metabolites from literature (L-data)^43–70^. On the L-data set, MetaReact again showed clear superiority, identifying 33.3% at Top-1 and reaching 60.0% accuracy at Top-3 (Figure 2D).

MetaReact’s advantage over rule-based systems stems from generalization beyond predefined transformation templates, while its edge over other Transformer models likely arises from the ReactSeq encoding that emphasizes reactive centers and a two-stage training paradigm combining large-scale USPTO pretraining with fine-tuning on curated drug metabolism reactions. Overall, these results demonstrate that MetaReact is robust when enzyme information is unavailable, offering broad generalization and practical value for early-stage metabolism assessment.

### 3.3 *Enzyme-completion*: substrate-based concurrent prediction of enzymes and metabolites

In the *enzyme-completion* setting, the model must jointly infer the catalyzing enzyme (family and subtype) and the resulting metabolite from a given substrate, which is an important scenario for mechanistic metabolism studies where both the enzyme and its products are unknown.

To evaluate MetaReact’s performance in this setting, we used an enzyme-annotated subset of our internal test set comprising 2,073 reactions. As shown in Figure 3A-B, MetaReact maintained high accuracy across both non-CYP and CYP classes. For non-CYP enzymes, it reached 93% Top-3 accuracy for UGTs and >70% for aldehyde dehydrogenases (ALDHs), carboxylesterases (CESs), and SULTs, and achieved 100% for rare families such as NATs and EPHXs. MetaReact considerably exceeded the class-prior baseline for low-frequency enzymes, indicating that its performance reflects genuine enzyme recognition rather than bias toward dominant classes. These results collectively demonstrate MetaReact’s robust ability to infer enzyme families and subtypes under class imbalance.

**Figure 3.**
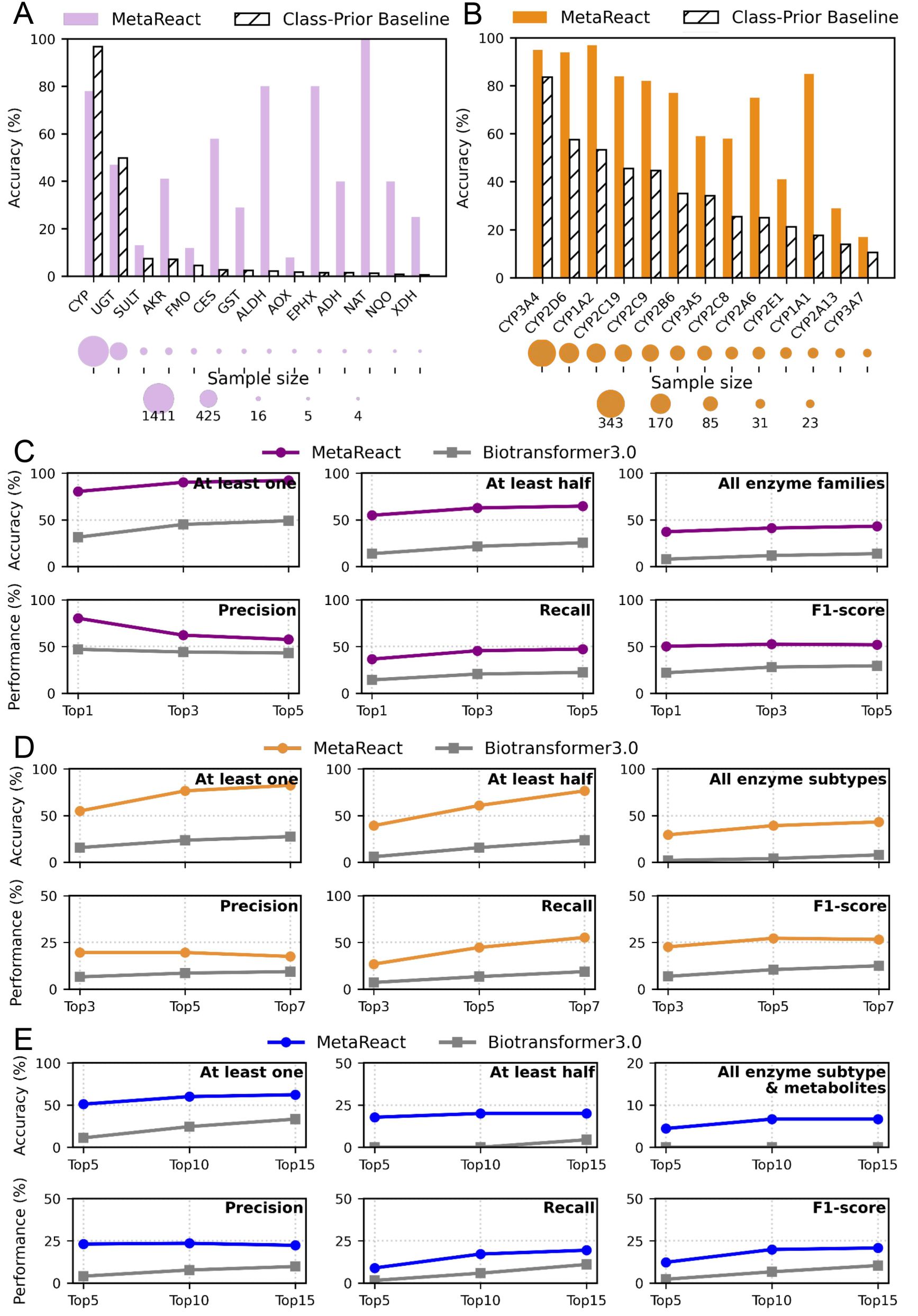
Comparison of prediction performance in the *enzyme-completion* setting. **(A, B)** Performance of MetaReact vs. random guessing on the internal test set, for (A) enzyme families and (B) CYP subtypes. Dot size reflects the sample size (number of reactions) for each enzyme family or subtype. (**C-E**) Comparison of MetaReact and BioTransformer3.0 on the external D-data benchmark. Panels show prediction performance at the (C) enzyme family, (D) enzyme subtype, (E) and enzyme subtype-metabolite pair levels, with precision, recall, and F1-score reported across Top-k thresholds. Accuracy is also evaluated under three coverage criteria: at least one, at least half, and all matched.

For further comparison with existing methods, we collected an external test se containing enzyme information (D-data set), by curating the latest entries from DrugBank v5.1.12^71^. It comprises 123 drugs and 198 corresponding metabolites. MetaReact consistently outperformed BioTransformer3.0 in all tasks (Figure 3C-E). At the family level, it achieved 80% Top-1 precision versus 50% for BioTransformer3.0, with an even larger advantage at the more challenging subtype level. MetaReact also outperformed BioTransformer3.0 when jointly predicting both the metabolizing enzyme subtype and the resulting metabolites.

Overall, MetaReact demonstrates strong concurrent prediction of both enzymes and metabolites, surpassing BioTransformer3.0 and performing robustly across frequent and rare enzyme classes. This highlights its value for real-world metabolism prediction, where enzyme information is often sparse, imbalanced, or unavailable.

### 3.4 Enzyme-conditioned: improved metabolite ranking and site-of-metabolism identification

In the *enzyme-conditioned* setting, the model is provided both the substrate and the catalyzing enzyme, enabling metabolite prediction under specified catalytic contexts. This scenario reflects common medicinal chemistry workflows, where enzyme involvement is experimentally established and chemists seek to anticipate likely metabolites as well as vulnerable SOMs.

As shown in Figure 4A, incorporating enzyme context significantly improved MetaReact’s accuracy on the internal test set, with Top-1 precision increasing from 28.8% to 46.2%, indicating that the model effectively leverages enzyme information to make context-aware predictions. We further evaluated MetaReact on the external D-data benchmark, comparing it with the rule-based tool CyProduct, which is restricted to nine major CYP450 isoforms. This evaluation tested whether incorporating enzyme context enables the model to generate more accurate rankings of likely products. Under matched enzyme conditions (Figure 4B), MetaReact achieved slightly higher Top-1 accuracy than CyProduct, and its advantage widened at broader cutoffs. At Top-2 and Top-3, MetaReact showed markedly higher recall, and at Top-10 and Top-20 it recovered more than 70% of reference metabolites, while CyProduct failed to generate predictions beyond Top-3. Beyond this comparison, MetaReact also supports both CYP and non-CYP enzymes.

**Figure 4.**
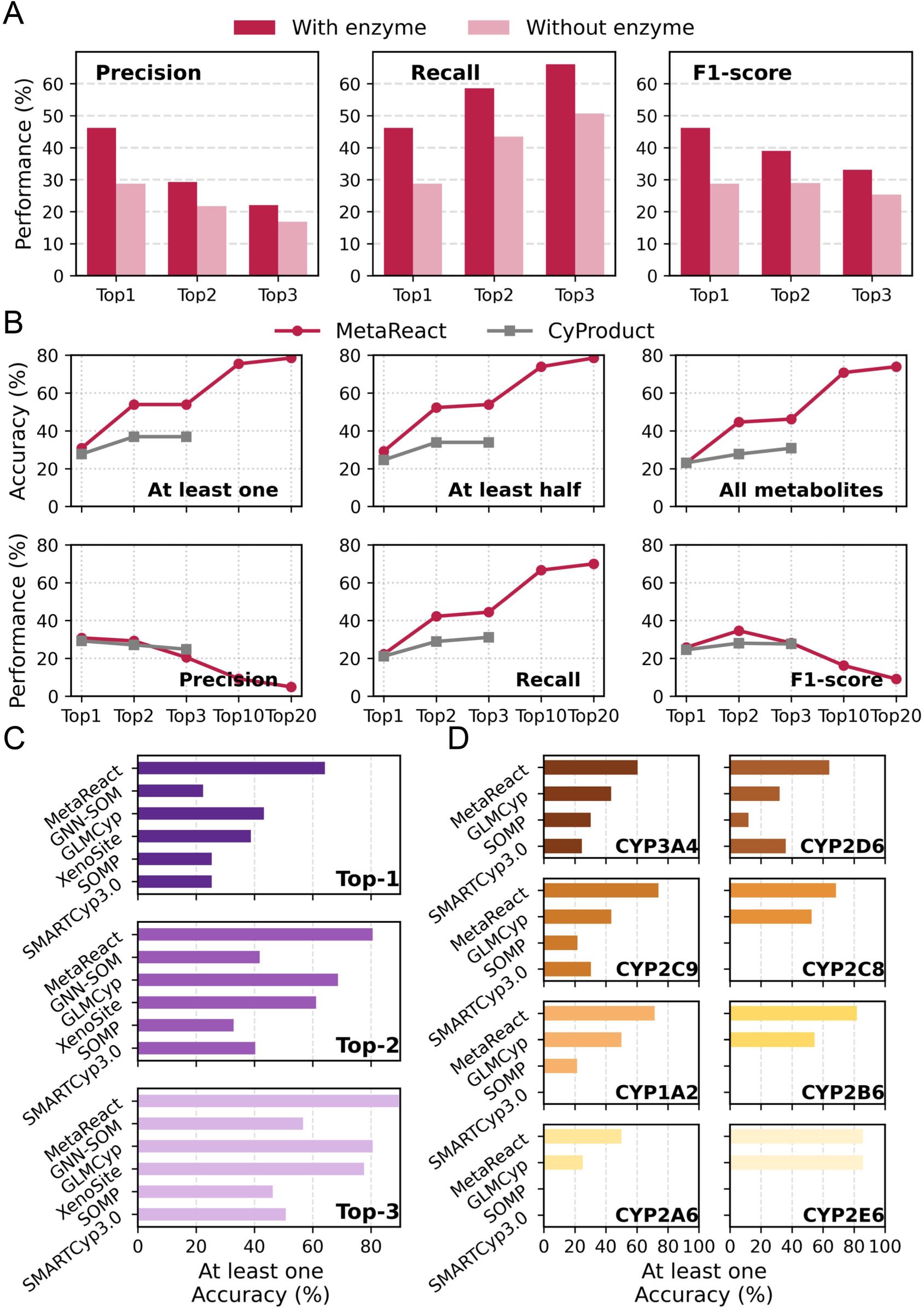
Performance of MetaReact in the *enzyme-conditioned* setting. **(A)** Effect of enzyme context on metabolite prediction in the internal test set. Performance was evaluated by precision, recall, and F1-score with and without enzyme subtype information. **(B)** Metabolite-prediction performance of MetaReact and the rule-based tool CyProduct under matched CYP450 isoform conditions. Accuracy, precision, recall, and F1-score are compared at multiple Top-k cutoffs. **(C-D)** SOM-prediction performance of MetaReact and baseline models at two levels: (C) CYP450 family level, showing Top-1, Top-2, and Top-3 accuracies, and (D) isoform level, showing Top-1 accuracy.

We further examined the prediction of SOMs, a task central to medicinal chemistry where chemists identify metabolically vulnerable positions to reduce liabilities and avoid toxic products. Evaluation was performed on an external SOM test set (357 annotated sites from 74 drugs), focused on CYP-mediated metabolism. At both the family level (Figure 4C) and isoform level (Figure 4D), MetaReact consistently achieved the highest accuracies, clearly outperforming all baselines. GLMCyp ranked second overall and was the only other method able to cover all eight isoforms, but its performance was still lower than MetaReact across cutoffs. Rainbow XenoSite and GNN-SOM showed reasonable accuracy at broader cutoffs but relied heavily on fixed auxiliary information, limiting adaptability to unseen contexts. SMARTCyp3.0 and SOMP performed competitively only within subsets of isoforms and fell behind elsewhere, reflecting the broader limitations of rule-based and descriptor-driven methods. By contrast, MetaReact employs a reaction-centric representation that captures atom- and bond-level transformations directly from molecular graphs, enabling broader enzyme coverage and stronger generalization beyond predefined rules.

Together, these results demonstrate that MetaReact effectively integrates enzyme specificity to guide both metabolite ranking and SOM identification, offering a unified and broadly applicable framework for metabolism prediction across CYP and non-CYP contexts.

### 3.5 Predictions in Challenging Scenarios

#### *Enzyme-Agnostic*: Substrate-Only Metabolite Prediction in Complex Cases

Accurate prediction of metabolites directly from substrates is essential for drug safety, pharmacological innovation, and public health, especially when catalyzing enzymes are unknown. Such challenges are exemplified by synthetic cannabinoids (SCs), which undergo rapid and extensive metabolism that hinders forensic detection^72^; plant-derived natural products (NPs), whose structurally diverse constituents lack clear metabolic characterization; and clinical drugs that still encounter unanticipated liabilities in humans^73^. These scenarios represent complex cases where predictive tools capable of capturing multi-step metabolic transformations are needed to complement experimental assays and support both research and regulatory decision-making.

##### Synthetic Cannabinoids (SCs)

MetaReact successfully predicted diverse metabolic pathways for SCs (Figure 5A). For 5F-EDMB-PICA, the model identified dealkylation and aliphatic hydroxylation with downstream transformations^74^, matching reported data. For hexahydrocannabinol (HHC), hydroxylation site predictions were consistent with experimental evidence.^51^ The model also captured the amide hydrolysis of 1cP-LSD, aligning with known profiles.^58^ These results underscore the model’s utility in supporting forensic screening and toxicology through early metabolite identification.

**Figure 5.**
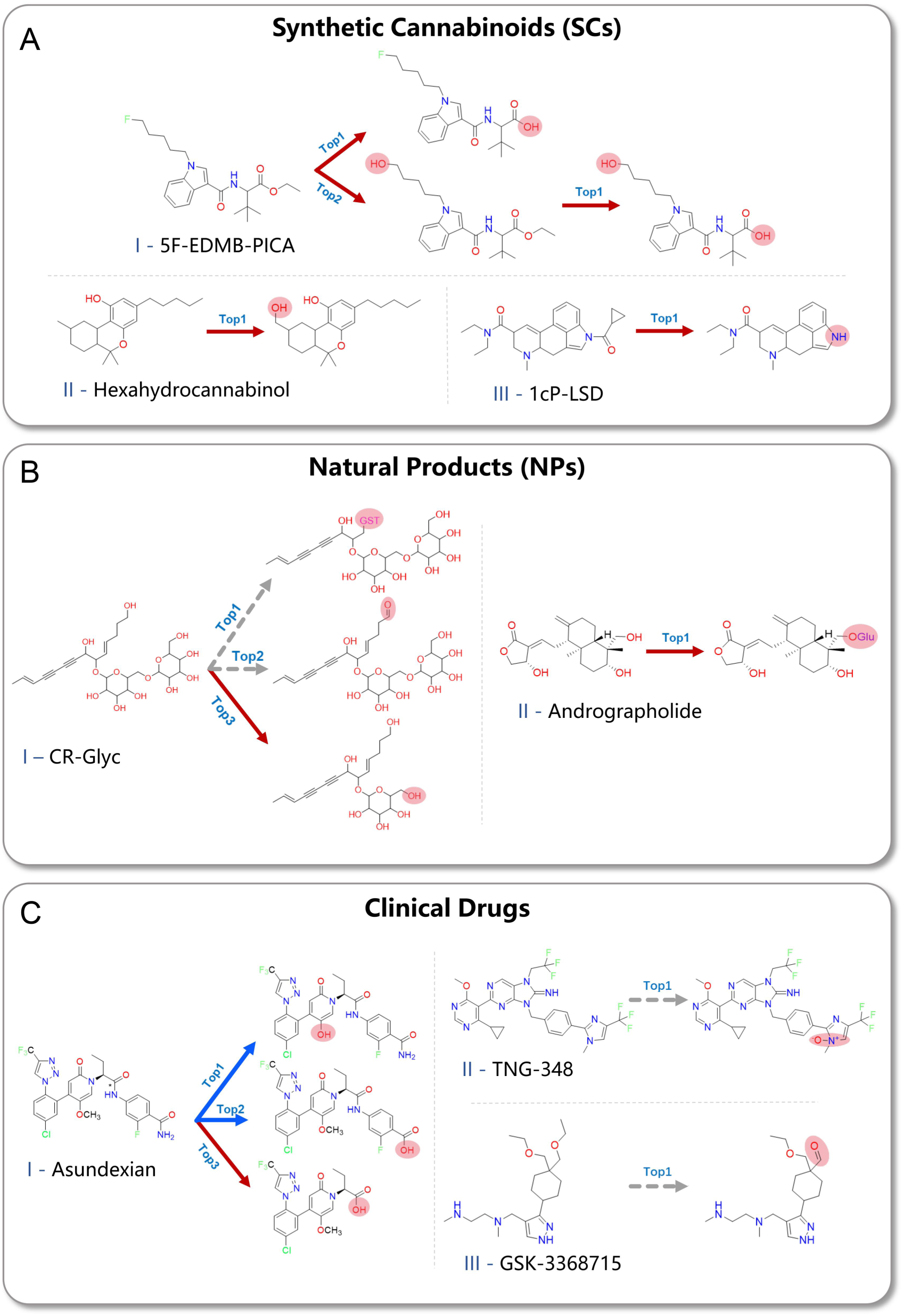
Case studies of metabolite prediction for challenging compound classes. Predicted metabolic pathways of **(A)** Synthetic Cannabinoids (SCs), **(B)** Natural Products (NPs) and **(C)** Clinical Drugs. Red and blue solid arrows point to experimentally confirmed major and minor metabolites, respectively, while grey dashed arrows indicate predicted metabolites that have not yet been experimentally validated.

##### Natural Products (NPs)

For CR-Glyc, a representative glycosylated constituent isolated from *Codonopsis Radix*, MetaReact predicted deglycosylation, which is typically mediated by gut microbiota^62^, and also suggested possible glutathione conjugation and oxidative metabolites (Figure 5B), although these require further experimental confirmation. For andrographolide, sulfonation and glucuronidation were correctly identified as major Phase II pathways, consistent with literature reports^75^. The predictions demonstrate the model’s potential to aid systematic NP metabolism research and modernization.

##### Clinical Drugs

MetaReact accurately captured both primary and secondary human metabolites across representative clinical and preclinical examples (Figure 5C). For Asundexian (Factor XIa inhibitor), the model correctly identified amide hydrolysis as the dominant metabolic route and further detected two minor pathways involving dealkylation and amide-to-acid conversion.^50^ To explore safety-related failures, we additionally predicted the potential metabolites of discontinued candidates. For TNG-348, a ubiquitin-specific protease 1 (USP1) inhibitor, the model predicted N-oxidation at a tertiary amine site, suggesting formation of reactive intermediates potentially responsible for the observed Grade 3-4 hepatotoxicity in Phase I/II trials (NCT06065059). For GSK-3368715, a protein arginine methyltransferase (PRMT) inhibitor, side-chain oxidation was predicted to yield electrophilic aldehyde intermediates, consistent with its Phase I termination due to safety concerns.^76^ Collectively, these results demonstrate MetaReact’s ability to not only predict major and minor human metabolites but also reveal mechanistic links between metabolic transformations and clinical safety risks.

#### Enzyme-Completion: AOX-Related Clinical Failures

Jointly identifying the catalyzing enzyme and its metabolites is critical for mechanistic metabolism studies and drug development, as overlooking enzyme involvement can lead to costly late-stage failures. AOXs exemplify this challenge: they are Phase I molybdenum-containing enzymes with pronounced species differences, being highly expressed in humans and cynomolgus monkeys but scarcely present in standard preclinical models such as mice, rats, and dogs. Furthermore, AOXs localize to the cytosol rather than the microsomal fraction commonly used in in vitro assays, making their activity difficult to evaluate. These features often cause AOX-mediated metabolism to be underestimated during preclinical studies, resulting in unexpected pharmacokinetic outcomes in clinical trials.

MetaReact achieved high accuracy in predicting AOX-related transformations. For example, Falnidamol failed in development due to low oral bioavailability from AOX1-mediated metabolism^77^; MetaReact ranked AOX1 third among predicted enzymes and correctly generated its primary metabolite. In another case, SGX523 caused renal toxicity due to insoluble AOX-derived metabolites^78^; MetaReact ranked AOX1 first and accurately predicted the toxic metabolite structure (Figure 6A). These results highlight the value of MetaReact in early pharmacokinetic profiling, enabling researchers to anticipate AOX metabolism and select animal models with AOX activity more comparable to humans (e.g., cynomolgus monkeys) to mitigate species-specific metabolic discrepancies and reduce clinical trial risk.

**Figure 6.**
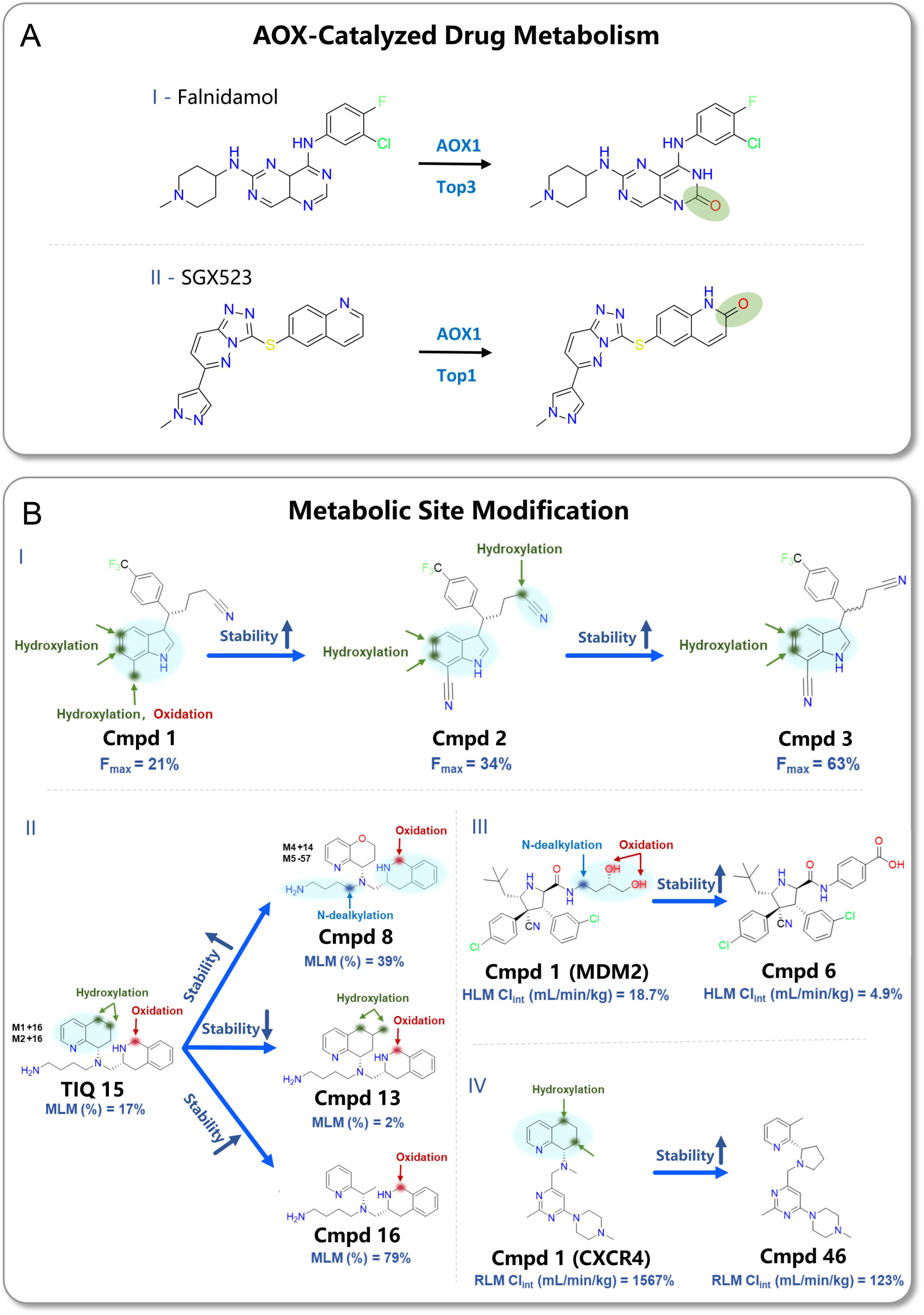
Prediction of AOX-catalyzed drug metabolism and metabolic site modification cases. **(A)** Falnidamol and SGX523, showing correct AOX1 ranking and metabolite prediction. **(B)** Representative site-directed modifications for CXCR4 antagonist TIQ-15 (I) and ZAK inhibitor ZAK-14 (II). MLM (%) refers to the percentage of compound remaining after 10 minutes of incubation with mouse liver microsomes. HLM Cl_int_ (mL/min/kg) represent the intrinsic clearance by human liver microsomes. The light-blue shaded fragment denotes the experimentally identified metabolic site region, within which metabolism is known to occur but without atom-level resolution. The arrow and highlighted atom(s) mark the specific metabolic site predicted by MetaReact.

#### Metabolic Site Modification Analysis

Identifying metabolic soft spots on a molecule is critical for rational compound optimization. By pinpointing metabolically vulnerable positions, medicinal chemists can modify or block these sites to improve stability, reduce clearance, and avoid the formation of toxic products. Accurate prediction of SOMs is therefore essential for designing drug candidates with enhanced pharmacological efficacy and optimized pharmacokinetic profiles.

MetaReact is inherently suited for SOM prediction because its reaction-aware representation, ReactSeq, explicitly encodes atomic and bond-level transformations, naturally capturing reaction centers where metabolic modifications occur. This enables the model to recognize structurally susceptible positions even without predefined enzyme information.

To demonstrate the capability of MetaReact in predicting metabolic hotspots, we analyzed four representative case studies, including C-X-C chemokine receptor type 4 (CXCR4) antagonists^79, 80^, mineralocorticoid receptor antagonists^7^, and murine double minute 2 homolog (MDM2) inhibitors (Figure 6B).^81^

In one example (Figure 6B-I), Bayer’s non-steroidal mineralocorticoid receptor antagonist Cmpd1 showed poor metabolic stability in hepatocytes. Experimental profiling identified predominant oxidation at the indole 5-/6-position and two-step oxidation of the methyl group.^7^ MetaReact accurately captured both the vulnerable sites and reaction types. The acid metabolite formed in the second oxidation step suggests possible involvement of ALDH in addition to CYP-mediated oxidation. This observation highlights that MetaReact does not rely on predefined enzyme families and can effectively operate in complex systems such as HLM, where multiple enzymes coexist. Subsequent analogs with a cyano substitution (Cmpd2) or truncated side chain (Cmpd3) displayed improved stability, and MetaReact correctly reflected the shifting metabolic liabilities, such as loss of reactivity at the α-carbon adjacent to the cyano group.

Beyond this series, MetaReact also highlighted oxidation and dealkylation liabilities in TIQ-15, Cmpd1 (MDM2), and Cmpd1 (CXCR4), as shown in Figure 6B-II, -III and -IV, consistent with experimental observations. It further reproduced how structural modifications affected stability, including incorporation of pyran rings, nitriles, or esters. In some cases, it even explained design failures, such as methylation of Cmpd13 introducing new oxidation sites that worsened clearance. Further implementation details are provided in the Supplementary Information Section B.1.

Overall, these cases illustrate that MetaReact not only predicts metabolic hotspots in line with experiments but also provides mechanistic insights into why certain modifications improve or reduce stability, thereby offering actionable guidance for medicinal chemists.

## 3 Conclusion and Discussion

Drug metabolism research seeks to understand enzymatic transformations that underlie drug efficacy and toxicity, thereby informing the design of safer and more effective therapies. Existing computational tools are often fragmented, constrained to specific enzyme families or rule-based heuristics, and rarely optimized for clinically critical tasks such as major metabolite prioritization.

Here we present MetaReact, a unified Transformer-based framework that integrates metabolite prediction, enzyme annotation, and site-of-metabolism identification in a single model. Its performance is driven by three innovations: the ReactSeq representation, which explicitly encodes atom- and bond-level transformations; a pretraining-fine-tuning strategy, which transfers general chemical reactivity knowledge to metabolism-specific data; and a prompt-guided task formulation, which allows one model to flexibly adapt to *enzyme-agnostic*, *enzyme-completion*, and *enzyme-conditioned* settings. Together, these designs enabled MetaReact to outperform state-of-the-art methods across diverse benchmarks.

Each setting aligns with distinct real-world applications. In the *enzyme-agnostic* mode, MetaReact successfully predicted metabolites of structurally complex substrates such as synthetic cannabinoids, traditional Chinese medicines, and clinical drugs. In the *enzyme-completion* mode, it identified aldehyde oxidase involvement in failed clinical compounds, highlighting hidden enzymatic liabilities. In the *enzyme-conditioned* mode, it provided actionable guidance for metabolic site modification, supporting rational optimization strategies.

Despite these advances, several challenges remain. Model accuracy is still limited by the coverage and balance of training data, especially for rare enzymes and multi-step pathways. ReactSeq focuses on local transformations but does not fully capture conformational dynamics or protein-ligand interactions. Moreover, real-world applications demand the ability to capture drug-drug interactions, such as competitive metabolism when multiple compounds share the same enzyme, and to achieve cross-species generalization from animal studies to human contexts where enzymatic activity differs substantially. Addressing these limitations will require tighter integration of structural biology, multi-omics perturbation data, and experimental validation.

In conclusion, MetaReact establishes a scalable, rule-free paradigm for human metabolism modeling. By unifying metabolite, enzyme, and SOM prediction and demonstrating broad applicability from complex substrates to clinical pipelines, it provides both methodological innovation and practical impact, laying the foundation for more comprehensive metabolism-aware drug discovery.

## 4 Methodology

### 2.1 Preparation of Metabolic Reaction Datasets

We curated a large-scale collection of enzymatic metabolic reactions from multiple public sources, including DrugBank^71^, MetXBioD^82^, SMPDB^83^, HumanCyc^84^, Recon3^85^, and the Accelrys Metabolite Database (AMD; now BIOVI)^86^. The collected reactions were curated by (1) removing known cofactors from multi-product reactions, and (2) mapping enzyme names to UniProt (UniProt ID)^87^ and NCBI Entrez Gene^88^, using gene symbols as standardized enzyme identifiers to harmonize data across sources. The resulting dataset contains 62,695 unique reactions across 15 enzyme families and 138 subtypes (Figure 7A). Substrate and product molecular weights span 30-900 Da, with most reactions involving drug-like molecules (Figure 7B-C).

**Figure 7.**
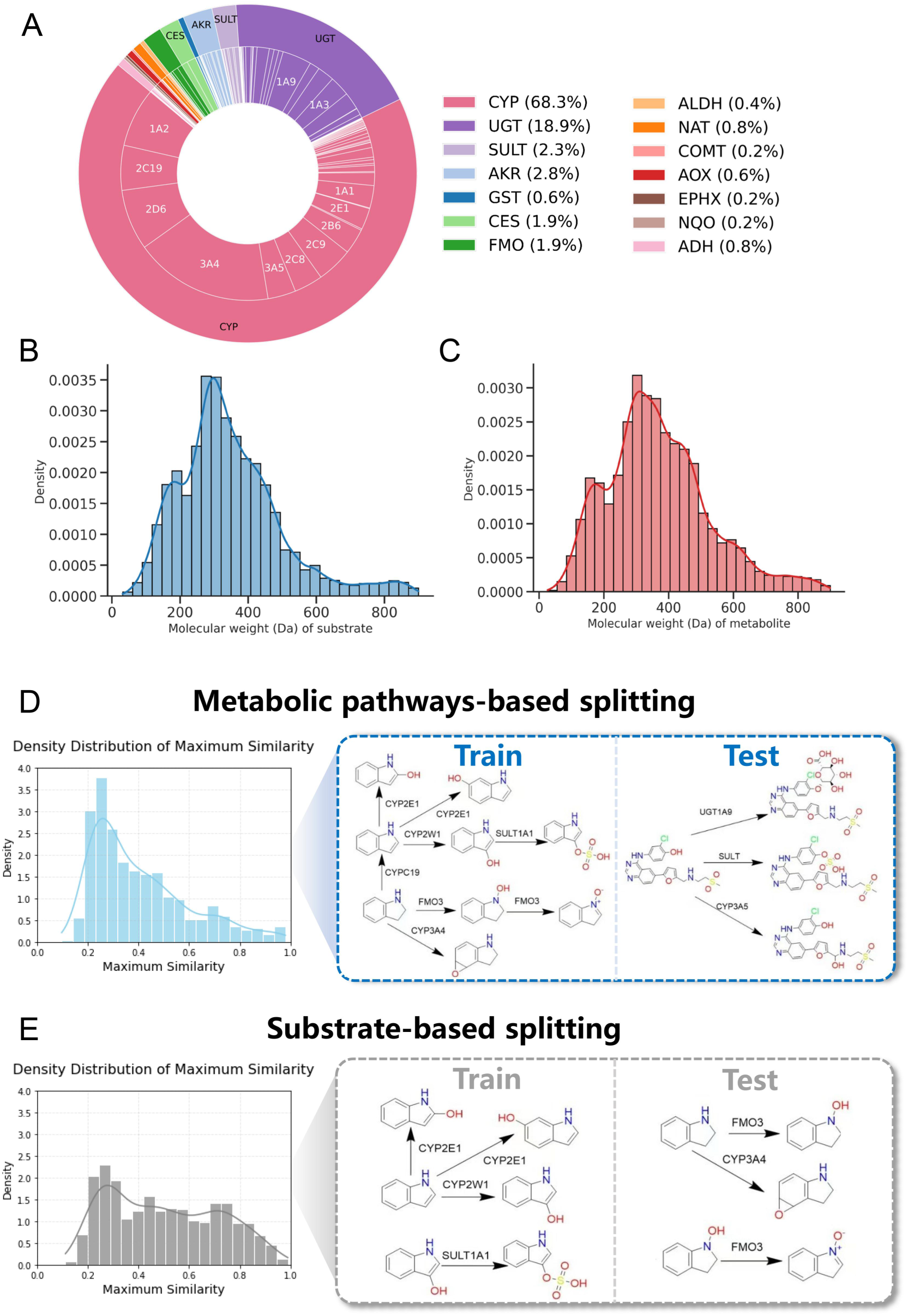
Composition of the curated dataset and splitting strategies. **(A)** Distribution of metabolizing enzymes by family and subtype. **(B-C)** Molecular weight distributions of substrates and metabolites. **(D-E)** Comparison of maximum Tanimoto similarity between training and test sets under different splitting schemes, with schematics illustrating pathway-based versus substrate-based partitioning.

To ensure strict independence between training and evaluation, the fine-tuning dataset was partitioned by metabolic pathways, a stricter strategy than the commonly used substrate-based split (Figure 7D-E). This approach ensures a higher degree of dissimilarity among the training, validation, and test sets, thereby mitigating the risk of data leakage due to molecular structure similarity. Specifically, we constructed a directed metabolic reaction graph using the NetworkX package^89^, where each node represents a reactant or product SMILES string, and edges represent enzyme-catalyzed transformations with directionality reflecting the reaction flow. Disconnected subgraphs within this metabolic network were extracted and saved as independent units. These subgraphs were then allocated to the training, validation, and internal test sets in an 8:1:1 ratio, based on the total number of reactions (i.e., edges) they contained. To preserve data independence, all reactions within a given subgraph were assigned exclusively to a single dataset split.

#### Pretraining Dataset from USPTO

We employed the USPTO chemical reaction dataset for model pretraining. The USPTO dataset, released by the United States Patent and Trademark Office, encompasses a comprehensive collection of patent records. We specifically used the USPTO-MIT subset, which consists of 479,035 curated chemical reactions extracted via text mining from patents issued between 1976 and September 2016.^42^ Since metabolic reaction prediction uses a single molecule as input, all reactions were represented in the reverse (retrosynthetic) direction to ensure consistency with this formulation.

#### Reaction Representation by ReactSeq

All reaction data were encoded using ReactSeq^41^, a reaction-centric molecular representation language we previously developed. ReactSeq systematically encodes retrosynthetic transformations by defining a set of molecular editing operations (MEOs), which explicitly describe atomic and bond-level changes occurring in a chemical reaction. Originally proposed for single-step retrosynthesis, ReactSeq has demonstrated substantial improvements in prediction accuracy for retrosynthetic planning tasks. The core design principle of ReactSeq is to represent a reaction as a sequence of discrete transformation steps, enabling stepwise reconstruction of one or more reactants from a given product (Figure 1B). This reaction-aware encoding provides a more intuitive and detailed view of chemical transformations and is fully interconvertible with standard SMILES notation. To handle cases where only major substrate-product pairs are known, ReactSeq inserts a placeholder token “Au” for unobserved leaving groups, which is removed post-prediction. Because ReactSeq explicitly represents the formation and breaking of chemical bonds, it inherently highlights the reactive centers of a molecule, making it particularly suitable for learning SOM information.

#### Augmentation Strategies for Training and Inference

During both pretraining and fine-tuning, we expanded the data by applying a 10-fold augmentation in which the initial atom ordering of SMILES strings was randomized, enabling more robust learning of reactant-product transformations. At inference, test-time augmentation (TTA) was performed using the same randomization strategy, generating 10 augmented inputs per reactant. For each input, beam search was used with a configurable beam width (beam_size) to generate 20 candidate products, yielding a total of 200 predictions per reactant. Final scores were assigned by integrating beam-search rankings with candidate frequencies across augmentations. Additional architectural choices, optimization settings, and scheduler details are provided in the Supplementary Information Section A.1 and Figure S1.

### 2.2 Model Architecture and Training

#### Transformer Backbone and Tokenization

MetaReact adopts a sequence-to-sequence Transformer with an encoder-decoder backbone. SMILES strings are encoded at the character level, while enzyme identifiers are added to the vocabulary as single, indivisible tokens. Missing enzyme information is represented by a placeholder token “ <blank>”, which is excluded from loss computation via masking. The tokenizer maps characters to integer indices that are subsequently embedded into high-dimensional vectors and processed by self-attention in the encoder and cross-attention in the decoder. All reactions are represented in ReactSeq to expose atom- and bond-level transformations. Details are provided in the in the Supplementary Information Section A.2.

#### Pretraining and Fine-tuning Pipeline

Training follows a transfer-learning pipeline. The model is first pretrained on ReactSeq-formatted reverse reactions derived from general organic chemistry for 500,000 epochs. The learned weights are then transferred and fine-tuned on the curated metabolic reaction dataset for 10,000 epochs, yielding the final MetaReact model. Model selection is based on validation performance.

### 2.3 Evaluation Framework Evaluation Metrics

Model performance was quantified by computing the Tanimoto similarity (based on ECFP4 fingerprints) between predicted and true metabolites, with a similarity score of 1 indicating a correct match. The performance metrics included accuracy, precision, recall, and F1-score. The specific calculation formulas are as follows:

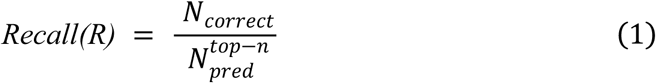

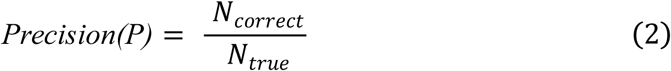

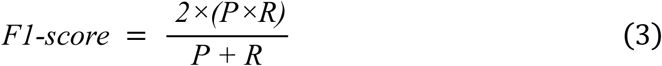

where N*_correct_* denotes the number of correctly predicted metabolites, 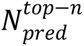 is the total number of metabolites predicted in the top-n results, and N*_true_* is total number of ground-truth metabolites.

Given that a substrate may yield multiple true metabolites, we further define three match-level evaluation metrics. *At least one metabolite* indicates at least one predicted metabolite exactly matches any of the true metabolites. *At least half metabolites* shows at least 50% of the ground-truth metabolites are recovered by the prediction. *All metabolites* means the predicted set of metabolites is exactly consistent with the reference set. Hereafter, accuracy refers to the *At least one metabolite* metric unless stated otherwise.

#### *Enzyme-agnostic* evaluation (MetaTrans dataset & L-data)

In the *enzyme-agnostic* setting, evaluation was performed on the MetaTrans dataset, introduced by Litsa et al.^38^, which contains 65 drugs and 179 experimentally validated metabolites, including 29 CYP450 substrates from the GLORY benchmark and 36 non-CYP substrates newly added in DrugBank v5.1.7.

In addition, we used a literature-derived dataset (L-data), manually curated from 28 peer-reviewed publications over the past two years (Table S3).^43–70^ Each entry in L-data pairs a drug with its manually annotated major metabolite, allowing evaluation of the model’s ability to prioritize clinically relevant outcomes. Full details of dataset composition and baseline configurations are provided in the Supplementary Information Section A.3.

#### *Enzyme-completion* evaluation (D-data & internal test set)

In the *enzyme-completion* setting, evaluation was performed on two datasets. The D-data benchmark contains 123 newly added drugs and their 198 corresponding metabolites curated from DrugBank version 5.1.12^71^ relative to version 5.1.10. Among these, 51 drugs have annotated enzyme information spanning six families and 19 subtypes (Figure S2). To extend enzyme coverage, we further incorporated 2,073 reactions with explicit enzyme annotations from the internal test dataset. This subset spans 14 enzyme families and 72 subtypes, including CYP, UGT, sulfotransferase (SULT), and AOX, with CYP450 and UGT accounting for ∼70% and ∼20% of reactions, respectively (Figure S3).

#### *Enzyme-conditioned* evaluation (D-data & internal test set)

In the *enzyme-conditioned* setting, evaluation was performed in two scenarios. First, we used 2,073 enzyme-annotated reactions from the internal test set to compare predictions with and without enzyme subtype information, thereby quantifying the benefit of enzyme context. Second, we benchmarked MetaReact against CyProduct^36^, a rule-based tool covering nine major CYP450 isoforms (CYP1A2, CYP2A6, CYP2B6, CYP2C8, CYP2C9, CYP2C19, CYP2D6, CYP2E1, CYP3A4). For fairness, MetaReact’s predictions were restricted to the same enzyme conditions; in the D-data benchmark, this corresponded to 38 drugs with 65 annotated reactions.

#### SOM prediction (SOM external test set)

SOM prediction aims to identify the specific atoms within a substrate that undergo enzymatic transformation. In this setting, evaluation was conducted using an external SOM test set comprising 357 enzyme-substrate-site entries from 74 drugs, constructed by combining D-data with CYP-related reactions from MetaTrans dataset. This dataset spans eight major CYP450 isoforms (CYP3A4, CYP2C9, CYP2D6, CYP2C8, CYP2B6, CYP2A6, CYP1A2, CYP2E1).

MetaReact was benchmarked against existing CYP450-focused SOM predictors, including SOMP^24^, SMARTCyp3.0^25^, Rainbow XenoSite^27^, GLMCyp^29^, and GNN-SOM^28^. Two evaluation settings were considered. At the CYP450 family level, the enzyme family was specified without distinguishing isoforms; GNN-SOM and Rainbow XenoSite were evaluated only in this setting as they do not support isoform-specific prediction. At the CYP450 isoform level, performance was assessed for eight representative isoforms, each with at least 20 reactions to ensure reliability (Figure S4). SMARTCyp3.0 and SOMP were tested on a subset of isoforms, whereas GLMCyp and MetaReact covered all eight.

## Data availability

The Drugbank data can be downloaded from https://go.drugbank.com/#. The MetXBioDB dataset is available at https://zenodo.org/records/4247792. The Recon3D dataset can be accessed at https://ngdc.cncb.ac.cn/databasecommons/database/id/6846. The HumanCyc dataset can be downloaded from the following link: https://maayanlab.cloud/Harmonizome/dataset/HumanCyc+Pathways. The SMPDB is available at https://smpdb.ca.

## Code availability

The MetaReact source code, trained model and implementation scripts are are publicly available at our GitHub repository: https://github.com/myzhengSIMM/MetaReact.

## Acknowledgements

This study was supported by the Strategic Priority Research Program of the Chinese Academy of Sciences (XDB0830000, China), National Natural Science Foundation of China (82204278, China; T2225002, China and 82273855, China), SIMM-SHUTCM Traditional Chinese Medicine Innovation Joint Research Program (E2G805H, China), Shanghai Municipal Science and Technology Major Project, National Key Research and Development Program of China (2023YFC2305904, China and 2022YFC3400504, China), Key Technologies R&D Program of Guangdong Province (2023B1111030004, China), Shanghai Post-doctoral Excellence Program (2024707, China), Postdoctoral Fellowship Program of CPSF (GZB20250838, China) and the Lingang Laboratory (LGL-8888-02, China).

We also acknowledge Shanghai Supercomputer Center for providing computing resources.

## Author Contributions

**Yitian Wang** and **Jingxin Rao**: Writing—Original draft preparation. **Wei Zhang**: Formal analysis. **Yuqi Shi**: Formal analysis. **Chuanlong Zeng**: Formal analysis. **Rongrong Cui**: Formal analysis. **Yinquan Wang**: Formal analysis. **Jiacheng Xiong**: Formal analysis and Supervision. **Xutong Li**: Formal analysis and Supervision. **Mingyue Zheng**: Conceptualization and Supervision.

