## Supplementary Information for "MetaReact: A Reaction-Aware Transformer for End-to-End Prediction of Drug Metabolism"

A. **Supplementary Methods**

**A.1 Data augmentation**

We employed a data augmentation strategy to improve model training. During training, both the pretraining reaction dataset and the metabolic reaction dataset were augmented by randomizing the initial atom ordering in SMILES strings. The corresponding aligned ReactSeq representations were then generated, yielding a 10-fold augmentation based on different atom index permutations. This procedure was designed to facilitate the learning of reactant–product transformations in ReactSeq.

At the inference stage, we adopted test-time augmentation (TTA). Specifically, the atom ordering of input SMILES was randomized to produce 10 augmented variants per reactant. Beam search, a heuristic search algorithm that retains the top-ranked candidates at each decoding step, was then applied. The beam width (b) controls the number of candidates preserved per step. For each augmented input, beam search generated 20 candidate products, ranked in descending order of their cumulative probabilities. The resulting ReactSeq predictions were converted into standardized metabolite SMILES, with invalid and duplicate strings removed. Figure S1 illustrates the TTA procedure, using an example with t = 3 and b = 5.

Given 10 augmented inputs per reactant and a beam width of 20, a total of 200 candidate products were obtained. To score these candidates, we combined their beam-search ranking positions with their frequencies of occurrence across augmentations, as formalized in the following equation:

$$\mathrm{Score}(output)=\sum_{n=1}^{t} \sum_{k=1}^{b} \frac{1}{k^{2}}$$

In this framework, *t* denotes the augmentation factor and *b* denotes the beam width. Predictions ranked higher in beam search received larger weights, while candidates appearing more frequently accumulated higher scores. The final predictions were then selected based on the overall scores in descending order.

**A.2 Model Architecture and Training Pipeline**

We constructed MetaReact based on a sequence-to-sequence Transformer architecture. The model is composed of an attention-based Transformer backbone comprising an encoder and a decoder. The encoder employs self-attention to process the input sequence and generate context-aware representations, with each layer capturing progressively abstract features of the molecular input. The decoder follows an autoregressive strategy, where each token is generated sequentially based on self-attention over the previously generated tokens. Additionally, cross-attention mechanisms allow the decoder to dynamically integrate contextual information from the encoder, enabling token-by-token generation of the metabolic output sequence.

In our implementation, we adopted the OpenNMT-py framework (version 3.2.0)^1^ with a character-level tokenizer, which encodes each character in a SMILES string as its corresponding ASCII value. Enzyme name abbreviations were also incorporated into the vocabulary. For entries lacking enzyme information, which represent the majority of the dataset, we inserted a special placeholder token ("<blank>") at the enzyme position. These padding tokens were ignored during model training. The final vocabulary size was 366. The tokenizer splits input strings into discrete tokens, which are then mapped to unique numerical indices through the vocabulary. These indices are passed into the Transformer model and embedded into high-dimensional vectors for further processing. The model architecture was configured with a word vector size (word_vec_size) and hidden size (hidden_size) of 256, six layers in both the encoder and decoder, eight attention heads, and a feed-forward dimension (transformer_ff) of 2048. Optimization was performed using the Adam optimizer with an initial learning rate of 1, coupled with the Noam learning rate scheduler (β₁ = 0.9, β₂ = 0.998, warm-up steps = 8000). During training, we set the batch size to 64 and applied a 30% dropout rate to mitigate overfitting. The batch type was set to sents, meaning batches were organized by sentence count. For model inference, beam search was employed with a configurable beam width (beam_size) and the number of returned hypotheses (n_best) to retrieve the top-ranked candidate sequences.

As to the training pipeline of MetaReact, we first pretrained the model on a reverse reaction dataset represented in ReactSeq format, derived from general organic chemical reactions. The learned weights were then transferred as initialization for the downstream metabolite prediction model. This pre-trained model was subsequently fine-tuned on a curated dataset of drug metabolism reactions, also represented using the ReactSeq scheme, resulting in the final MetaReact model. During pre-training, the model was trained for 500,000 epochs with model checkpoints saved every 1,000 epochs. Fine-tuning was conducted over 10,000 epochs, with model parameters saved every 100 epochs and evaluation on the validation set performed every 200 epochs. The final model weights used for external testing were selected based on the best validation performance.

Task-specific prompts were applied to guide the model in handling distinct prediction scenarios. The enzyme-agnostic prompt enables metabolite prediction without providing any enzyme information. The enzyme-completion setting allows the model to simultaneously predict both the enzyme and the corresponding metabolite. The enzyme-conditional prompt is used when the enzyme is known and the task is to predict the associated metabolite. This unified prompting strategy allows a single model architecture to flexibly address multiple tasks without requiring task-specific retraining, significantly enhancing both the versatility and practical usability of the model.

**A.3 Baseline methods and evaluation protocols**

Rule-based approaches – SyGMa^2^ applies Phase I and II transformation rules curated from 6,187 reactions in the MDL Metabolite Database and ranks metabolites by empirical likelihood scores. GLORYx^3^ integrates the random forest-based FAME3^4^ model for SOM prediction with rule-based metabolite generation. BioTransformer^5^ combines expert-derived rules with machine learning components (e.g., random forests) for small-molecule metabolism prediction.

Transformer-based approaches – MetaTrans^6^ and MetaPredictor^7^ adopt Transformer architectures and employ beam search to generate and rank candidate metabolites.

For ranking-enabled models (GLORYx, SyGMa, MetaReact, MetaPredictor, MetaTrans), predictions were sorted by model scores, and evaluation was performed at multiple cutoffs (Top-5, Top-10, Top-13, and Top-20). At each cutoff, accuracy was assessed using three reaction coverage metrics: (i) at least one correct metabolite recovered; (ii) at least half of the reference metabolites recovered; and (iii) all reference metabolites recovered. In addition, recall and precision were computed at each cutoff to evaluate retrieval and ranking quality.

For BioTransformer, which lacks a native ranking mechanism, predictions are returned as an unranked set, and the number of predicted metabolites varies by input. Across the 65 substrates in M-data, the average output size is 13 candidates. To enable fair comparison with ranked models, BioTransformer’s accuracy and recall were calculated using this average output size, aligning it with the Top-13 evaluation point.

In addition to M-data benchmarking, we conducted an external evaluation using a benchmark dataset (L-data) specifically curated to assess model performance in ranking major metabolites. Each entry in L-data contains a single, manually annotated major metabolite for a given drug. Model performance was evaluated using recall at Top-1, Top-3, and Top-5 thresholds. Since BioTransformer does not generate ranked outputs and GLORYx was temporarily inaccessible, the comparison was restricted to MetaReact, MetaPredictor3 MetaTrans, and SyGMa.

B. **Supplementary Results**

**B.1 SOM cases**

TIQ-15, a CXCR4 antagonist, exhibited poor metabolic stability in mouse liver microsomes (MLM Cl_int_ = 15%). LC–MS/MS analysis revealed rapid oxidation on the 5,6,7,8-tetrahydroquinoline (THQ) ring.^8^ MetaReact accurately predicted THQ hydroxylation consistent with experiment and further suggested potential oxidative ring opening on the 1,2,3,4-tetrahydroisoquinoline (THIQ) moiety. Introduction of an oxygen atom into THQ (Cmpd 8) improved stability (MLM Cl_int_ = 39%) but shifted metabolism to THIQ, forming oxidation/dehydrogenation and N-dealkylation metabolites, both correctly captured by the model. Methyl substitution (Cmpd 13) reduced stability (MLM Cl_int_ = 2%) due to newly exposed THQ and side-chain oxidation sites, whereas THQ ring opening (Cmpd 16) enhanced stability (MLM Cl_int_ = 79%) by eliminating all THQ liabilities, leaving only THIQ oxidation predicted.

Cmpd1 (MDM2), as an MDM2 inhibitor, showed high clearance in human liver microsomes (18.7 mL/min/kg).^9^ The model predicted oxidation on the diol hydroxyl groups and N-dealkylation on the side chain, consistent with experimental confirmation that the diol side chain was a major metabolic site. Substituting the diol with a terephthalic acid moiety yielded Cmpd6, reducing clearance to 4.9 mL/min/kg, with the model predicting effective blocking of the side-chain reactive sites.

Poor oral bioavailability has been attributed to rapid hepatic microsomal clearance. Cmpd1 (CXCR4) showed high intrinsic clearance in rat liver microsomes (1567 mL/min/kg), with the THQ moiety primarily responsible.^10^ The model predicted multiple hydroxylation reactions on the THQ ring. Ring opening of the THQ moiety produced Cmpd46, decreasing intrinsic clearance to 123 mL/min/kg, with model predictions confirming that the original THQ reactive sites were blocked, accounting for improved metabolic stability.

C. **Supplementary Figures and Tables**

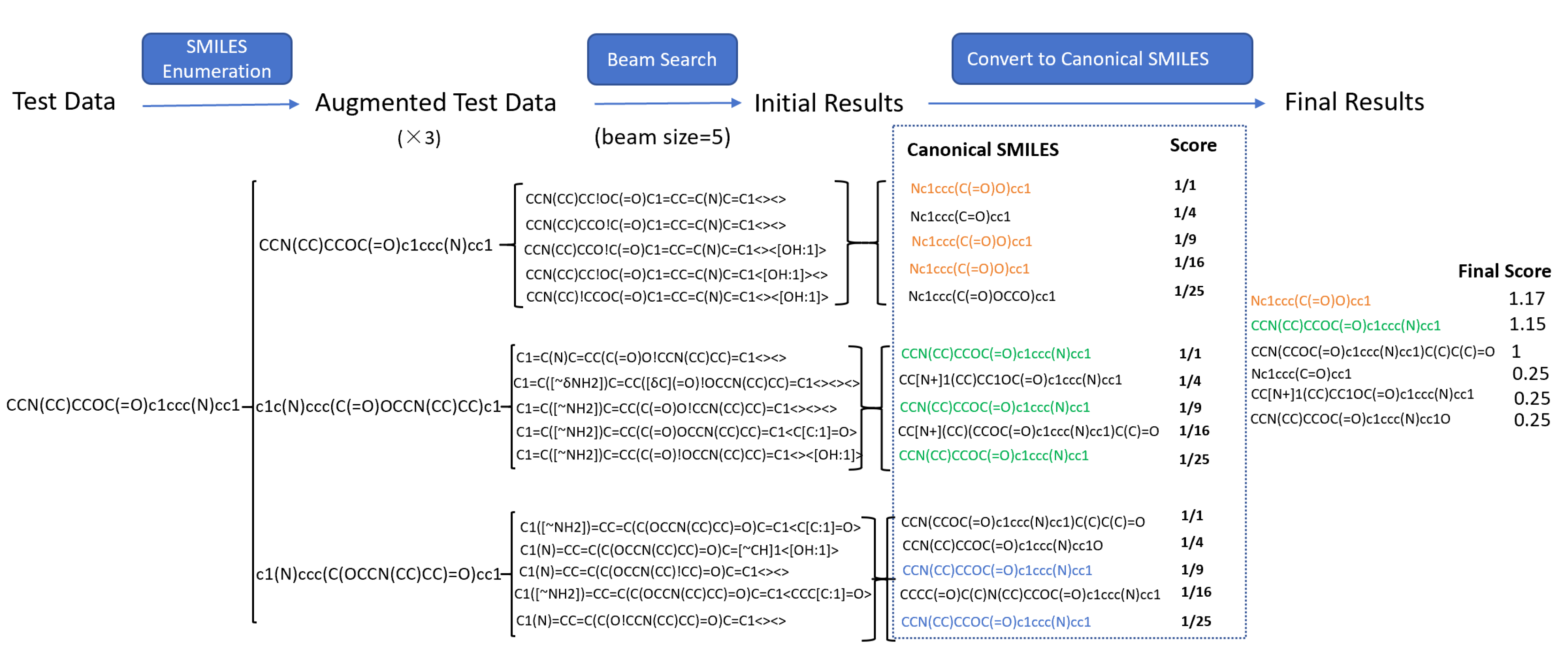

Figure S 1. Illustration of TTA strategy

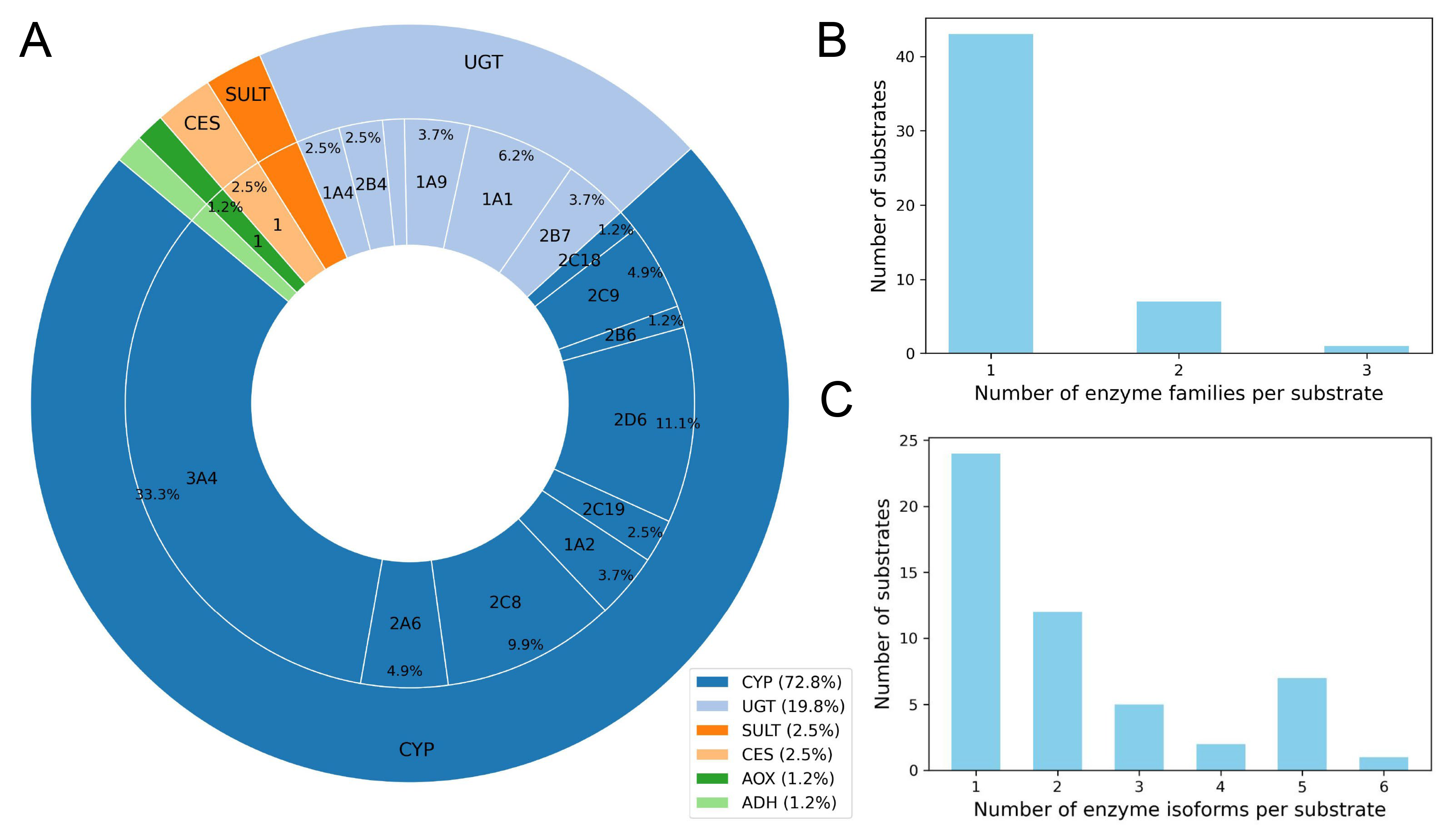

Figure S 2. Types and distribution of enzymes in the D-data dataset

**(A)** Distribution of enzyme families and subtypes. **(B–C)** Number of enzyme families (B) and subtypes (C) associated with each substrate.

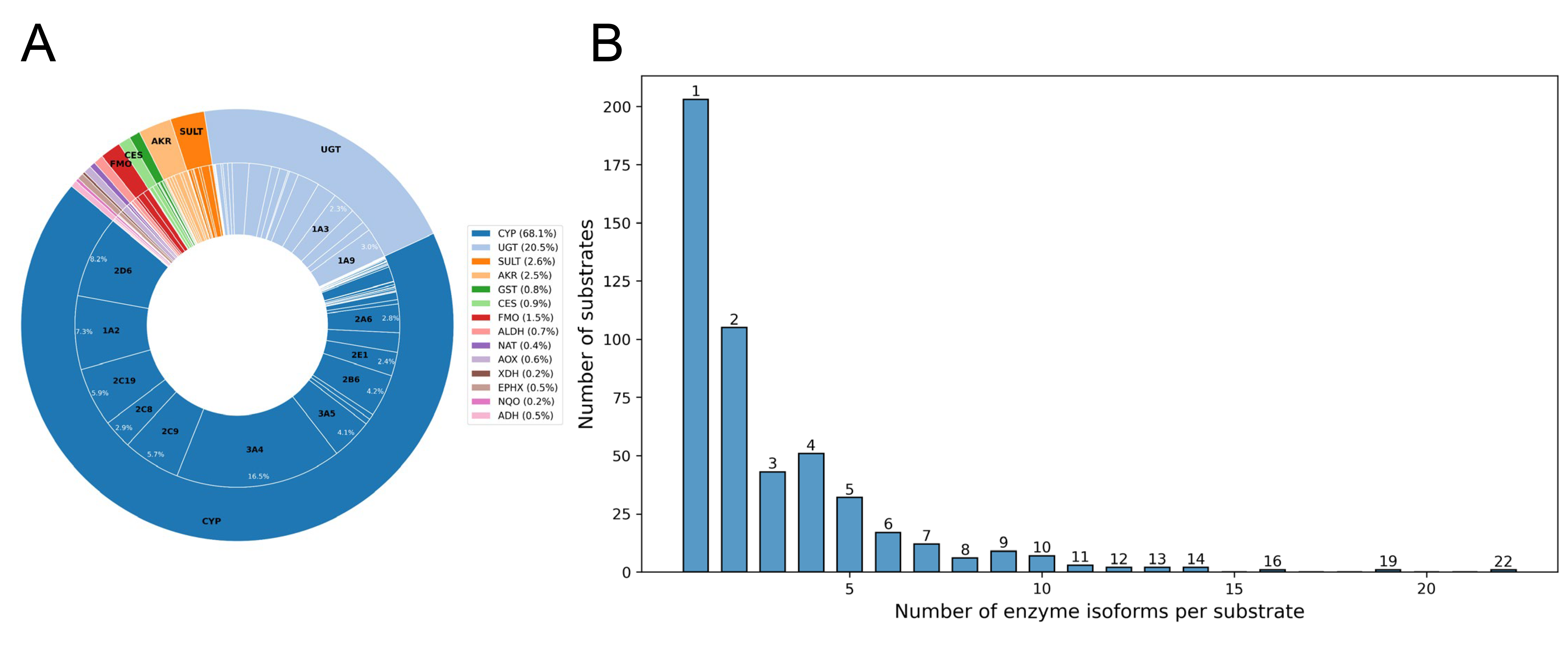

Figure S 3. Types and distribution of enzymes in the internal test set

**(A)** Distribution of enzyme families and subtypes. **(B)** Number of enzyme subtypes associated with each substrate.

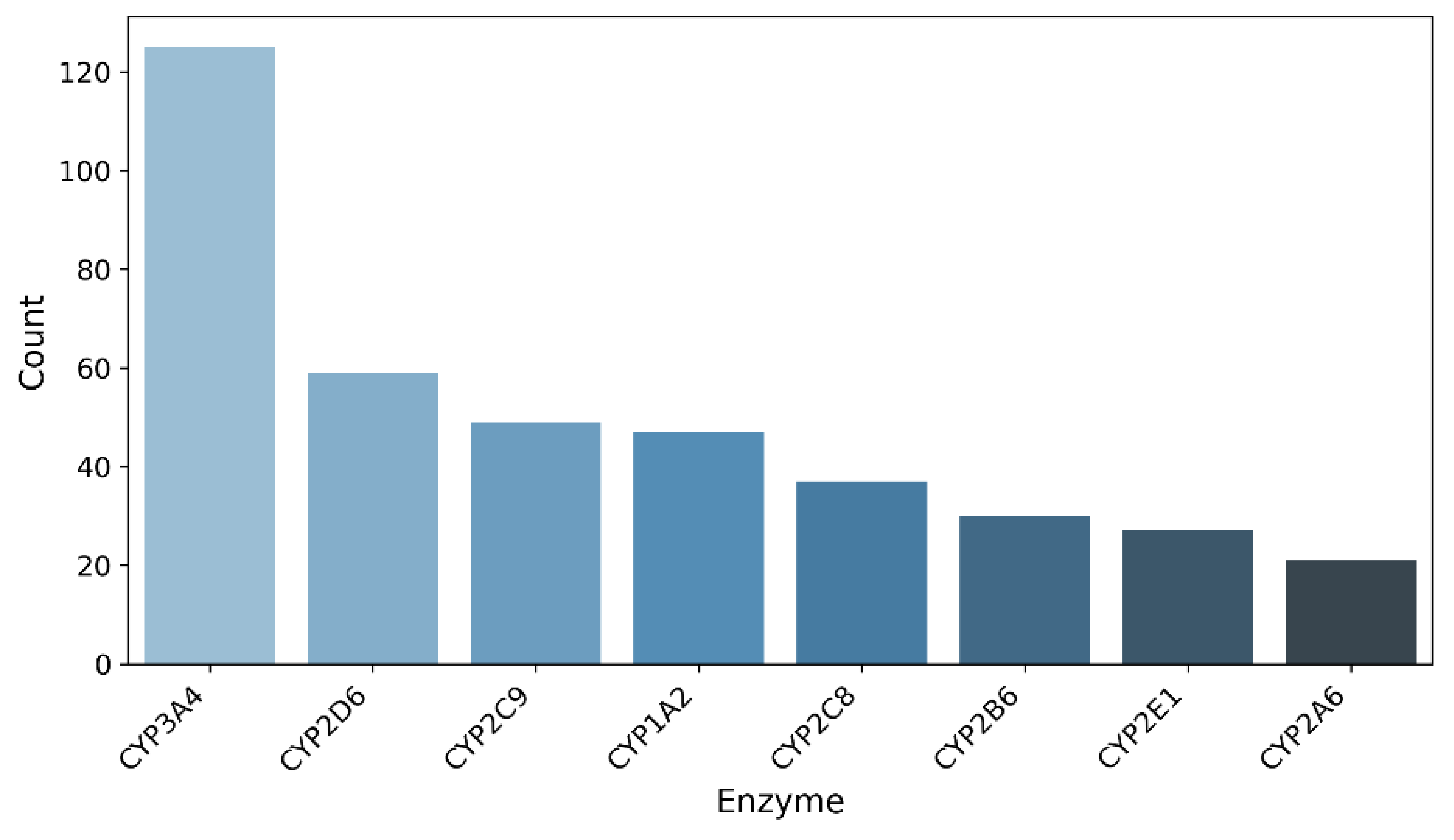

Figure S 4. Distribution of eight representative CYP enzyme isoforms in the SOM external test set

Table S 1. Comparative performance of different models on M-data by enzyme family

| Model/Dataset | Oxidatases | UGT | Sulfotransferases | Other Transferases | Hydrolases | Unspecified | Total |
| --- | --- | --- | --- | --- | --- | --- | --- |
| Dataset | 100 | 17 | 4 | 8 | 10 | 80 | 219 |
| MetaReact | 90 | 8 | 3 | 3 | 6 | 24 | 134 |
| MetaPredictor | 80 | 7 | 1 | 1 | 6 | 22 | 117 |
| MetaTrans | 70 | 7 | 3 | 2 | 4 | 23 | 109 |
| GLORYx | 70 | 8 | 3 | 1 | 4 | 22 | 108 |
| SyGMa | 80 | 8 | 2 | 0 | 5 | 20 | 115 |
| BioTransformer | 81 | 7 | 2 | 0 | 5 | 20 | 115 |

Table S 2. Comparative performance of different models on M-data in the *enzyme-agnostic* setting

| Top n | Model | At least one metabolite | At least half metabolites | All metabolites | Precision | Recall |
| --- | --- | --- | --- | --- | --- | --- |
| Top5 | MetaReact | **89.2%** | **72.3%** | **41.5%** | **32.3%** | **58.7%** |
|  | MetaPredictor | 87.7% | 70.8% | 35.4% | 30.2% | 54.7% |
|  | MetaTrans | 80.0% | 61.5% | 29.2% | 23.5% | 42.5% |
|  | GLORYx | 64.6% | 35.4% | 16.9% | 16.6% | 30.2% |
|  | SyGMa | 72.3% | 55.4% | 29.2% | 23.4% | 42.4% |
| Top10 | MetaReact | **95.4%** | **84.6%** | **55.4%** | **19.5%** | **70.9%** |
|  | MetaPredictor | 92.3% | 78.5% | 43.1% | 17.1% | 62.0% |
|  | MetaTrans | 95.4% | 80.0% | 44.6% | 15.0% | 57.5% |
|  | GLORYx | 80.0% | 64.6% | 27.7% | 14.3% | 51.9% |
|  | SyGMa | 87.7% | 75.4% | 43.1% | 16.2% | 58.7% |
| Top13 | MetaReact | **96.9%** | **90.8%** | **60.0%** | **15.9%** | **74.9%** |
|  | MetaPredictor | 93.8% | 83.1% | 44.6% | 11.2% | 65.4% |
|  | MetaTrans | 95.4% | 81.5% | 46.2% | 12.0% | 60.9% |
|  | GLORYx | 86.2% | 76.9% | 41.5% | 12.8% | 60.3% |
|  | SyGMa | 89.2% | 78.5% | 44.6% | 13.6% | 64.2% |
|  | BioTransformer | 87.7% | 78.5% | 44.6% | 13.5% | 64.2% |
| Top20 | MetaReact | **96.9%** | **95.4%** | **63.1%** | **11.0%** | **79.3%** |
|  | MetaPredictor | 95.4% | 86.2% | 50.8% | 9.5% | 68.7% |
|  | MetaTrans | 96.9% | 86.2% | 46.2% | 8.7% | 64.8% |
|  | GLORYx | 92.3% | 86.2% | 52.3% | 10.5% | 73.7% |
|  | SyGMa | 90.8% | 84.6% | 49.2% | 9.9% | 70.9% |

Table S 3. Overview of the L-data dataset

| Drug Name | Substrate SMILES | Major Metabolite SMILES | Exposure (%) | Source DOI |
| --- | --- | --- | --- | --- |
| Simnotrelvir (SIM0417) | CC(C)(C)[C@H](NC(=O)C(F)(F)F)C(=O)N1CC2(C[C@H]1C(=O)N[C@H](C#N)C[C@@H]1CCNC1=O)SCCS2 | CC(C)(C)[C@H](NC(=O)C(F)(F)F)C(=O)N1CC2(C[C@H]1C(=O)N[C@@H](C[C@@H]1CCNC1=O)C(=O)O)SCCS2 | 12.70 | <https://doi.org/10.1038/s41401-024-01393-7> |
| Asundexian | CC[C@@H](C(=O)Nc1ccc(C(N)=O)c(F)c1)n1cc(OC)c(-c2cc(Cl)ccc2-n2cc(C(F)(F)F)nn2)cc1=O | CC[C@@H](C(=O)O)n1cc(OC)c(-c2cc(Cl)ccc2-n2cc(C(F)(F)F)nn2)cc1=O | 47 | <https://doi.org/10.1007/s13318-023-00838-4> |
| Hexahydrocannabinol | CCCCCc1cc(O)c2c(c1)OC(C)(C)C1CCC(C)CC21 | CCCCCc1cc(O)c2c(c1)OC(C)(C)C1CCC(CO)CC21 | Not reported | <https://doi.org/10.1093/jat/bkad079> |
| HY-17542 | CC(=O)Nc1ccc(C)c(C(=O)N[C@H](C)c2cccc3ccccc23)c1 | Cc1ccc(N)cc1C(=O)N[C@H](C)c1cccc2ccccc12 | Not reported | <https://doi.org/10.3389/fphar.2023.1067408> |
| GRL0617 | Cc1ccc(N)cc1C(=O)N[C@H](C)c1cccc2ccccc12 | Cc1cc(O)c(N)cc1C(=O)N[C@H](C)c1cccc2ccccc12 | Not reported | <https://doi.org/10.3389/fphar.2023.1067408> |
| Brepocitinib | Cn1cc(Nc2nccc(N3C[C@H]4CC[C@@H](C3)N4C(=O)[C@@H]3CC3(F)F)n2)cn1 | Nc1nccc(N2C[C@H]3CC[C@@H](C2)N3C(=O)[C@@H]2CC2(F)F)n1 | Not reported | <https://doi.org/10.1124/dmd.124.001750> |
| Mobocertinib | C=CC(=O)Nc1cc(Nc2ncc(C(=O)OC(C)C)c(-c3cn(C)c4ccccc34)n2)c(OC)cc1N(C)CCN(C)C | C=CC(=O)Nc1cc(Nc2ncc(C(=O)OC(C)C)c(-c3cn(C)c4ccccc34)n2)c(OC)cc1N(C)CCNC | 12.70 | <https://doi.org/10.1124/dmd.124.001841> |
| Clotrimazole | Cc1ccc([C@@H]2O[C@H](CO)[C@@H](O)[C@H](O)[C@H]2O)cc1Cc1ccc(-c2ccc(F)cc2)s1 | Cc1ccc([C@@H]2O[C@H](CO)[C@@H](O)[C@H](OC3O[C@H](C(=O)O)[C@@H](O)[C@H](O)[C@H]3O)[C@H]2O)cc1Cc1ccc(-c2ccc(F)cc2)s1 | Not reported | <https://doi.org/10.1124/dmd.124.001812> |
| OSI-930 | O=C(Nc1ccc(OC(F)(F)F)cc1)c1sccc1NCc1ccnc2ccccc12 | O=C(Nc1ccc(OC(F)(F)F)cc1)c1sccc1NCc1cc(=O)[nH]c2ccccc12 | Not reported | <https://doi.org/10.1124/dmd.124.001802> |
| Capivasertib | NC1(C(=O)N[C@@H](CCO)c2ccc(Cl)cc2)CCN(c2ncnc3[nH]ccc23)CC1 | NC1(C(=O)N[C@@H](CCOC2O[C@H](C(=O)O)[C@@H](O)[C@H](O)[C@H]2O)c2ccc(Cl)cc2)CCN(c2ncnc3[nH]ccc23)CC1 | 28.20 | <https://doi.org/10.1124/dmd.124.001636> |
| Inh 1 | CCOc1ccc(NC(=S)N(CCO)Cc2cc3cc(C)cc(C)c3[nH]c2=O)cc1 | Cc1cc(C)c2[nH]c(=O)c(CN3CCOC3=N)cc2c1 | Not reported | <https://doi.org/10.1080/00498254.2024.2357765> |
| Cmpd15 | COc1ccc2c(Oc3ccc(C4(C(=O)NO)CCOCC4)cc3)ccnc2n1 | COc1ccc2c(Oc3ccc(C4(C(=O)NOC5O[C@H](C(=O)O)[C@@H](O)[C@H](O)[C@H]5O)CCOCC4)cc3)ccnc2n1 | 88.22 | <https://doi.org/10.1016/j.ejmech.2024.116853> |
| Lobetyolinin | C/C=C/C#CC#CC(O)C(/C=C/CCCO)OC1OC(COC2OC(CO)C(O)C(O)C2O)C(O)C(O)C1O | C/C=C/C#CC#CC(O)C(/C=C/CCCO)OC1OC(CO)C(O)C(O)C1O | Not reported | <https://doi.org/10.1016/j.jpba.2024.116339> |
| VU6032423 | Cc1ncc(S(=O)(=O)N2CCC(c3cn4ncnc4cc3Cl)CC2)s1 | O=S(=O)(c1cnc(CO)s1)N1CCC(c2cn3ncnc3cc2Cl)CC1 | 23.80 | <https://pubs.acs.org/doi/10.1021/acs.jmedchem.4c01193> |
| SHPL-49 | COc1ccc(CCCCOC2OC(CO)[C@@H](O)[C@H](O)[C@H]2O)cc1 | OCC1OC(OCCCCc2ccc(O)cc2)[C@H](O)[C@@H](O)[C@@H]1O | Not reported | <https://doi.org/10.1016/j.jpba.2024.116314> |
| Scutellarin | O=C(O)C1OC(Oc2cc3oc(-c4ccc(O)cc4)cc(=O)c3c(O)c2O)C(O)C(O)C1O | O=c1cc(-c2ccc(O)cc2)oc2cc(O)c(O)c(O)c12 | Not reported | <https://doi.org/10.1016/j.jpba.2024.116325> |
| Cmpd21 | CN1CCC[C@@H](OC(=O)[C@](O)(c2ccccc2)c2cccc(OCCCNC(=O)c3ccc(CNC[C@H](O)c4ccc(O)c5[nH]c(=O)ccc45)cc3)c2)C1 | O=C(NCCCOc1cccc([C@](O)(C(=O)O)c2ccccc2)c1)c1ccc(CNC[C@H](O)c2ccc(O)c3[nH]c(=O)ccc23)cc1 | 32 | <https://pubs.acs.org/doi/10.1021/acs.jmedchem.4c00298> |
| EC-5026 | CC[C@H](C)C(=O)N1CCC(NC(=O)Nc2ccc(OC(F)(F)F)c(F)c2)CC1 | CC[C@@H](CO)C(=O)N1CCC(NC(=O)Nc2ccc(OC(F)(F)F)c(F)c2)CC1 | 69.49 | <https://doi.org/10.1016/j.jpba.2024.116116> |
| Cmpd9a | CN(C(=O)Cc1cccs1)[C@@H](Cc1c[nH]c2ccccc12)C(=O)N[C@@](C)(Cc1ccccc1)C(=O)N[C@@H](CCCNC(=N)N)C(=O)N1CCC[C@H]1C(=O)N[C@@H](CCCNC(=N)N)C(N)=O | CN(C(=O)Cc1cccs1)[C@@H](Cc1c[nH]c2ccccc12)C(=O)N[C@@](C)(Cc1ccccc1)C(=O)N[C@@H](CCCNC(=N)N)C(=O)N1CCC[C@H]1C(=O)N[C@@H](CCCNC(=N)N)C(=O)O | Not reported | <https://pubs.acs.org/doi/10.1021/acsmedchemlett.4c00091> |
| 1cP-LSD | CCN(CC)C(=O)C1C=C2c3cccc4c3c(cn4C(=O)C3CC3)CC2N(C)C1 | CCN(CC)C(=O)C1C=C2c3cccc4[nH]cc(c34)CC2N(C)C1 | Not reported | <https://doi.org/10.1016/j.jpba.2024.116187> |
| VX-548 | COc1c([C@H]2C(C(=O)Nc3ccnc(C(N)=O)c3)O[C@@](C)(C(F)(F)F)[C@H]2C)ccc(F)c1F | C[C@H]1[C@@H](c2ccc(F)c(F)c2O)C(C(=O)Nc2ccnc(C(N)=O)c2)O[C@@]1(C)C(F)(F)F | 90.71 | [https://doi.org/10.1002/bdd.2387Digital Object Identifier (DOI)](https://doi.org/10.1002/bdd.2387) |
| 4-AcO-DET | CCN(CC)CCc1c[nH]c2cccc(OC(C)=O)c12 | CCN(CC)CCc1c[nH]c2cccc(O)c12 | Not reported | <https://doi.org/10.1016/j.jpba.2024.116187> |
| Cmpd7 | N#CC1CN(S(=O)(=O)c2cccc(C(=O)N3CCC[C@@H]3C(=O)NCc3ccc(C(F)(F)F)cc3)c2)C1 | NS(=O)(=O)c1cccc(C(=O)N2CCC[C@@H]2C(=O)NCc2ccc(C(F)(F)F)cc2)c1 | Not reported | <https://pubs.acs.org/doi/10.1021/acs.jmedchem.3c02413> |
| GS-5734 | CCC(CC)COC(=O)[C@H](C)N[P@](=O)(OC[C@H]1O[C@@](C#N)(c2ccc3c(N)ncnn23)[C@H](O)[C@@H]1O)Oc1ccccc1 | C[C@H](N[P@](=O)(OC[C@H]1O[C@@](C#N)(c2ccc3c(N)ncnn23)[C@H](O)[C@@H]1O)Oc1ccccc1)C(=O)O | Not reported | <https://pubs.acs.org/doi/10.1021/acs.jmedchem.4c00234> |
| CC-220 | O=C1CC[C@@H](N2Cc3c(OCc4ccc(CN5CCOCC5)cc4)cccc3C2=O)C(=O)N1 | O=C1CC[C@@H](N2Cc3c(OCc4ccc(CN5CCOCC5=O)cc4)cccc3C2=O)C(=O)N1 | 14 | <https://doi.org/10.1007/s13318-024-00886-4> |
| MDMA | CNC(C)Cc1ccc2c(c1)OCO2 | CC(N)Cc1ccc2c(c1)OCO2 | Not reported | <https://doi.org/10.1016/j.jpba.2024.116076> |
| Cmpd2 | Cn1cnc2ncn(Cc3nc([C@@H]4CO[C@@H](c5ccc(Cl)cc5)C4)no3)c(=O)c21 | Cn1cnc2[nH]c(=O)n(Cc3nc([C@@H]4CO[C@@H](c5ccc(Cl)cc5)C4)no3)c(=O)c21 | 43.40 | <https://pubs.acs.org/doi/10.1021/acs.jmedchem.3c02121> |
| Platycodin D | CC1OC(OC2C(OC(=O)[C@]34CCC(C)(C)CC3C3=CCC5[C@@]6(C)C[C@H](O)[C@H](OC7OC(CO)C(O)C(O)C7O)C(CO)(CO)C6CC[C@@]5(C)[C@]3(C)C[C@H]4O)OCC(O)C2O)C(O)C(O)C1OC1OCC(O)C(OC2OCC(O)C2(O)CO)C1O | CC1OC(OC2C(OC(=O)[C@]34CCC(C)(C)CC3C3=CCC5[C@@]6(C)C[C@H](O)[C@H](O)C(CO)(CO)C6CC[C@@]5(C)[C@]3(C)C[C@H]4O)OCC(O)C2O)C(O)C(O)C1OC1OCC(O)C(OC2OCC(O)C2(O)CO)C1O | Not reported | <https://doi.org/10.1016/j.jpba.2024.116016> |
| Soticlestat | O=C(c1cccnc1-c1ccncc1)N1CCC(O)(Cc2ccccc2)CC1 | O=C(O)[C@H]1OC(OC2(Cc3ccccc3)CCN(C(=O)c3cccnc3-c3ccncc3)CC2)[C@H](O)[C@@H](O)[C@@H]1O | 85.90 | <https://doi.org/10.1111/bcp.15917> |
| Alpibectir | O=C(CCC(F)(F)F)N1CCC2(CC1)CC(C(F)(F)F)=NO2 | O=C(O)C1OC(OC2(Cc3ccccc3)CCN(C(=O)c3cccnc3-c3ccncc3)CC2)[C@H](O)C(O)[C@@H]1O | 55 | <https://doi.org/10.1124/dmd.124.001562> |
